# *Macrobdella decora*: Old World Leech Gut Microbial Community Structure Conserved in a New World Leech

**DOI:** 10.1101/687418

**Authors:** Emily Ann McClure, Michael C. Nelson, Amy Lin, Joerg Graf

**Affiliations:** Department of Molecular and Cell Biology, University of Connecticut, Storrs, Connecticut, U.S.A.; Institute for Systems Genomics, University of Connecticut, Storrs, Connecticut, U.S.A.

## Abstract

Leeches are found in terrestrial, aquatic, and marine habitats on all continents. Sanguivorous leeches have been used in medicine for millennia. Modern scientific uses include studies of neurons, anticoagulants, and gut microbial symbioses. *Hirudo verbana*, the European medicinal leech, maintains a gut community dominated by two bacterial symbionts, *Aeromonas veronii* and *Mucinivorans hirudinis*, which sometimes account for as much as 97% of the total crop microbiota. The highly simplified gut anatomy and microbiome of *H. verbana* make it an excellent model organism for studying gut microbial dynamics. The North American medicinal leech, *Macrobdella decora,* is a hirudinid leech native to Canada and the northern U.S.A. In this study we show that *M. decora* symbiont communities are very similar to those in *H. verbana.* This similarity allowed for an extensive study in which wild caught animals were sampled to determine effects of geographic separation, time of collection, and feeding on the microbiome. Through 16S V4 rRNA deep sequencing we show that: i) the *M. decora* gut and bladder microbial communities are distinct, ii) the *M. decora* gut community is affected by feeding and long periods of starvation, and iii) geographic separation does not appear to affect the overall gut microbial community structure. We propose that *M. decora* is a replacement for *H. verbana* for studies of wild-caught animals and offer evidence for the conservation of annelid symbionts. Successful culturing and comparison of dominant symbionts from *M. decora* and *H. verbena* will provide the ability to assess host-symbiont co-evolution in future work.

**IMPORTANCE:** Building evidence implicates the gut microbiome in regulating animal digestion, nutritional acquisition, immune regulation, development, and even mood regulation. Because of the difficulty of assigning causative relationships in complex gut microbiomes a simplified model for testing hypotheses is necessary. Previous research in *Hirudo verbana* has suggested this animal as a highly simplified and tractable animal model of gut symbioses. Our data show that *Macrobdella decora* may work just as well as *H. verbana* without the drawback of being an endangered organism and with the added convenience of easy access to field-caught specimens. The similarity of the microbial community structure of species from two different continents reveals the highly-conserved nature of the microbial symbionts in sanguivorous leeches and confirms the medicinal leech as a highly simplified, natural animal model in which to study gut symbioses.

## INTRODUCTION

Leeches are a diverse animal group capable of surviving in freshwater, marine, and terrestrial environments. They are found on all continents and oceans on planet Earth (1, 2). Records of humans applying leeches medicinally survive from populations as far back as ancient Egypt (3, 4), resulting in the name medicinal leech, *Hirudo medicinalis* Linnaeus, 1758. As our understanding of hirundinid taxonomy improved, *Hirudo medicinalis* was subdivided into additional species including *H.verbana* Carena 1820 and *H.orientalis* Utevsky & Trontelj, 2005 (5, 6). Since 2004 in the United States, only *H. medicinalis* and *H. verbana* are approved for use as a medical device and must be shipped from suppliers in Europe (5). Although it shares the same common name, the North American medicinal leech, *Macrobdella decora*, was rarely used for blood letting. No mechanical or pharmaceutical product has yet been able to replicate the reduction of venous congestion achieved by the medical application *of Hirudo* leeches (briefly reviewed in (7)). This results in a continued need for medicinal leeches and a better understanding of their biology.

As approved medical devices, the natural feeding habits of leeches are exploited to reduce venous congestion and improve blood circulation in affected patients. To make the most out of unpredictable encounters with prey, hirudinid leeches consume up to five times their body weight in one feeding and can go up to 6-12 months between feedings (1). After initial attachment, the leech stimulates blood flow in the prey by secreting vasodilators and a number of anticoagulant peptides including hirudin and orthologues (8-11). The gastrointestinal system of *H. verbana* is highly simplified and consists of a pharynx, crop, and intestinum (1, 7). Once consumed, excess ions and water are rapidly removed from the blood meal to form a highly viscous intraluminal fluid (ILF) in the crop (7, 12). ILF remains in the crop over long periods of time before it slowly passes into the intestinum where it is digested (7).

The gut microbiome of medicinal leeches is especially simple when compared to common mammalian gastrointestinal models. Previous research described an ILF microbiota in *H. verbana* dominated by *Aeromonas veronii* and *Mucinivorans hirudinis* (9, 13-17). Subsequent studies revealed the presence of additional *Aeromonas* spp in *H. verbana* and other hirudinid leeches (18-20). *Clostridial* species have also been detected in culture-independent studies of the ILF of *H. verbana* (14, 21) and *H. orientalis* (20). Several functions for the dominant crop microbiota have been proposed including providing essential nutrients to the host (9, 22, 23), preventing other bacteria from colonizing, inhibiting putrefaction of the ILF (24), aiding the host’s immune responses (25, 26), or initiating the digestion of erythrocytes (27).

In most animals, the diversity of the microbiome increases along the length of the digestive tract and similar findings have been reported for *Hirudo* spp. The intestinum contains Alpha-, Gamma-, and Delta-proteobacteria, *Fusobacteria*, *Firmicutes,* and Bacteroidetes as well as *Aeromonas* and *Mucinivorans* (14). A number of closely related hirudiniform leech species have also tested positive for *Aeromonas* and Bacteroidetes from the digestive tract (8, 19, 20). The composition of the microbial community among the different *Hirudo* species studied is similar in both crop and intestinum (14, 17, 19, 20).

Unlike humans but similar to other annelids (28, 29) the bladder of *H. verbana* is colonized by a number of microbial species (30). Sequence analysis and fluorescence *in situ* hybridization micrographs of the *H. verbana* bladder show a stratified community consisting of *Ochrobactrum, Bdellovibrio, Niabella,* and *Sphingobacterium* (30). The difference in bladder microbiome when compared to the ILF or intestinum indicates that *H. verbana* microbiomes are body site specific and suggests a selection process that regulates the composition of these communities.

Much less is known about the gut microbiome of *M. decora.* Prior studies attempting to characterize the *M. decora* gut microbiome relied on aerobic culturing methods or sequencing total ILF DNA with primers specific for *Aeromonas* or Bacteroidetes symbionts. These studies revealed that the *Aeromonas* species associated with *M. decora* was *A. jandaei* (31) and the Bacteroidetes species was most similar to uncultured and unidentified species in a clade with *Rikenella, Mucinivorans,* and *Alistipes* (8). In this study, we strived for a more complete understanding of the *M. decora* gut microbiome through culture-independent 16S rRNA V4 deep sequencing with confirmation by fluorescence *in situ* hybridization (FISH). This procedure was also performed on *H. verbana* to compare microbial communities between the two host species using identical techniques.

In this work we describe the microbiota of the ILF, bladder, and intestinum in wild and laboratory-maintained specimens of the North American medicinal leech, *Macrobdella decora*. Comparison of core and common microbial operational taxonomic units (OTUs) from *M. decora* to those of the well-described *H. verbana* provides insights about the level of conservation of the microbiomes between distantly-related and geographically-isolated sanguivorous leeches.

## RESULTS

### Leech Organs Contain Distinct Microbial Communities

The microbial communities of three organs from two leech species (*Hirudo verbana* and *Macrobdella decora*) were analyzed in this study: the intestinum, crop, and bladder (Table S1). Due to their small size, for the intestinum and bladder the entire organ was homogenized, while for the crop only the ILF was collected. Total DNA was extracted from the samples and the V4 region of the 16S rRNA gene was amplified, sequenced using an Illumina MiSeq, and analyzed to determine the community composition using Qiime 1.9 .1 (32) and R 3.6.0. Of all parameters tested, the leech host species had the greatest effect on microbiome composition (PERMANOVA: F=316.77, R^2^=0.53, p=0.001) (Figure 1A). This difference diminished when performing the same analysis at a higher taxonomic level. For example, when the PERMANOVA was performed at the order level (88% sequence identity) the variation accounted for by host species still had the greatest effect on microbiome composition, but that effect decreased to only 25% (PERMANOVA: F=95.79, R^2^=0.25, p=0.001) (Figure 1B). This suggests that although the specific genera within the microbial community have changed between leech hosts, the general physiological functions performed by the microbiome have likely been conserved. An additional 47% of the variation between samples was not accounted for by host leech species alone and other factors are important in determining the community composition.

**Figure 1.**
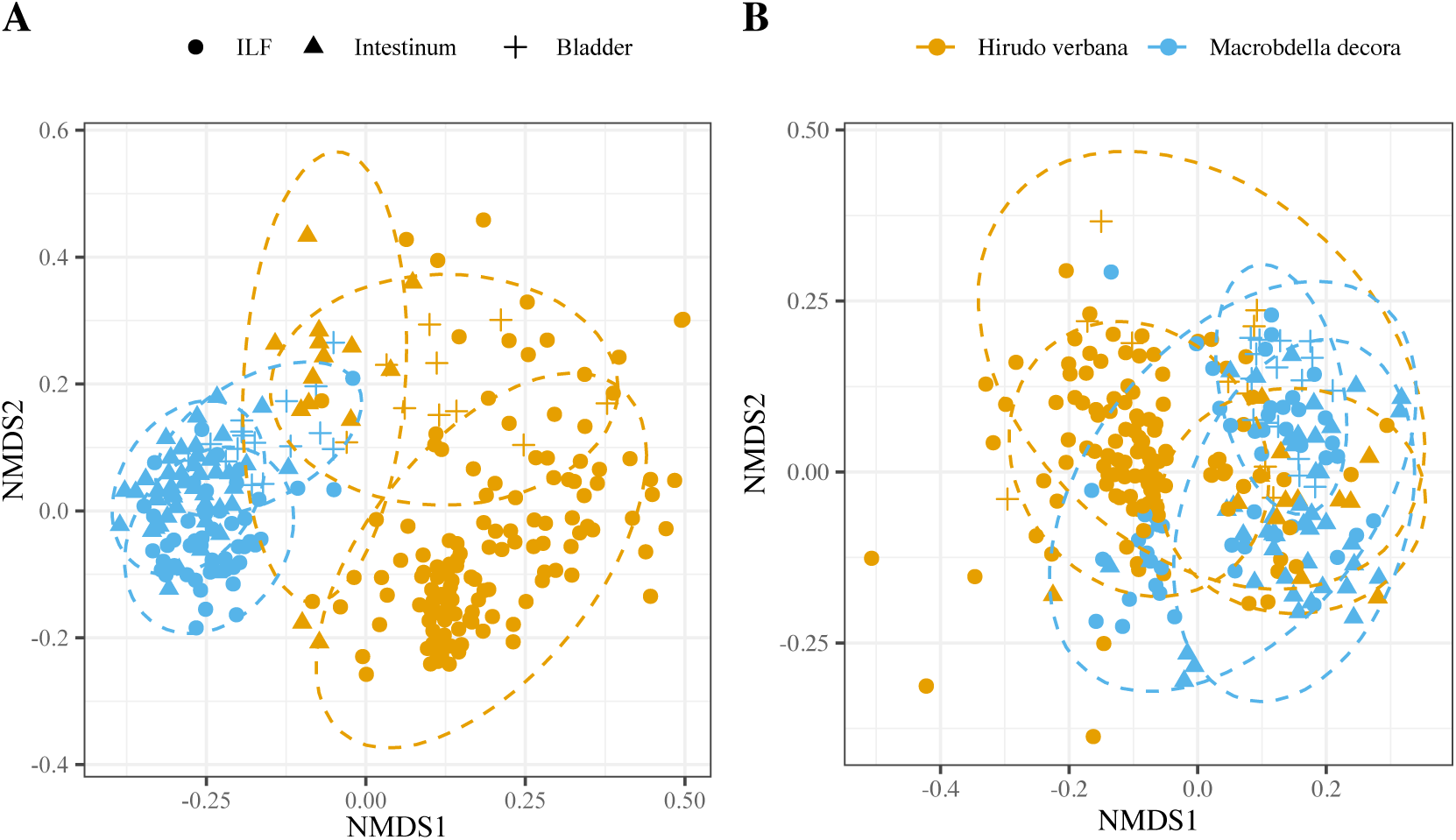
Unifrac-calculated NMDS plot of leech-associated microbiota showing that the microbiota is host species-specific at the genus level, but not at higher taxonomic levels. Two leech species were sampled (*Hirudo verbana* = orange and *Macrobdella decora* = cyan) at three organ sites (ILF = circle, intestinum = triangle, bladder = cross). A) When OTUs were calculated at 97% sequence similarity (∼Genus), 53% of the total variation between samples is described by separating the two leech host species. B) When OTUs were calculated at 88% sequence similarity (∼Order), only 25% of the total variation between samples is described by separating the two leech host species. Ellipses were calculated at 95% confidence.

In *H. verbana* samples, the sampled organ had the greatest effect on microbiome composition (PERMANOVA: F=57.56, R^2^=0.29, p=0.001) with feeding in the laboratory accounting for 18% of variation (PERMANOVA: F=3.90, R^2^=0.18, p=0.001), the supplier of *H. verbana* animals accounting for 5% of variation (PERMANOVA: F=19.71, R^2^=0.05, p=0.001), and the shipment date of *H. verbana* accounting for 7% of variation (PERMANOVA: F=6.63, R^2^=0.07, p=0.001). This confirms previous results that the bladder and digestive tract have very different microbiomes (14, 30) the microbial composition changes as a result of feeding (15). In addition, these data suggests that leech suppliers and date of shipment affect the observed ILF microbiome of *H. verbana*.

For this study, we obtained the *H. verbana* from two suppliers: one located in Germany that sold field-caught animals and one in France that sold farm-bred animals. Treatments before shipping, including prolonged starvation time to reduce abundance of human pathogens and potentially feeding animals antibiotic-contaminated blood (33), are additional sources of influence on microbiome variation accounted for but not uniquely discernable within the confounding influences clustered under the ‘supplier’ variable (18). Laboratory feeding in conjunction with sampled organ or animal source accounted for an additional 7% of variation (PERMANOVA: F=3.82, R^2^=0.03, p=0.001 and PERMANOVA: F=3.94, R^2^=0.04, p=0.001 respectively). This left ∼35% of variation unaccounted for by these four variables.

In *M. decora*, the sampled organ had the greatest effect on microbiome composition (PERMANOVA: F=26.47, R^2^=0.19, p=0.001) with the month of animal collection accounting for 17% of variation (PERMANOVA: F=9.23, R^2^=0.17, p=0.001), feeding in the laboratory accounting for 6% of variation (PERMANOVA: F=2.85, R^2^=0.06, p=0.001), and animal source accounting for only 5% of variation (PERMANOVA: F=4.37, R^2^=0.05, p=0.001). This mirrors the results from *H. verbana* in that the bladder and digestive tract have very different microbiomes, a change in microbial composition occurs as a result of feeding, and animal collection site and time also affect the observed ILF microbiome of *M. decora*.

Some variables additionally exhibited statistical significance when evaluated as a group. Sampled organ in conjunction with the month that the animals were collected accounted for an additional 15% of variation (PERMANOVA: F=5.16, R^2^=0.15, p=0.001). The animal collection month in conjunction with feeding accounted for a final 4% of variation (PERMANOVA: F=4.10, R^2^=0.04, p=0.001). This left ∼34% of variation unaccounted for by these four variables. Although the microbial differences between organs and after feeding were expected from previous studies of the *Hirudo* microbiome (14, 15, 19, 20, 30), it was surprising to discover the large role that collection month played in affecting the *M. decora* microbiome.

### Deep Sequencing of *H. verbana* ILF Microbiome Reveals Rare Community Members

16S rRNA V4 deep sequencing of the *H. verbana* ILF identified the same two dominant taxa as reported previously from 454 pyrosequncing of the 16S rRNA V6 region (21), 16S rRNA V3-V4 deep sequencing (17), and 16S rRNA clone library analysis (14) (Table 1). The ILF microbiota of *H. verbana* has been described as being dominated by *Mucinivorans* and *Aeromonas* with occasional Clostridial spp (14, 21). In the 36 animals tested, *Mucinivorans* and *Aeromonas* together accounted for 30.6 – 99.8% (median = 81.8%) of the sequences from the ILF (Table 1) with *Proteiniclasticum* making up an additional 0 – 63.4% (median when present = 10.8%, present in 69% of animals) and *Fusobacterium* making up an additional 0 – 28.3% (median when present = 11.5%, present in 47% of animals) (Table S2). The *Proteiniclasticum* OTU was previously reported in the ILF 16S rRNA gene clone libraries of *Hirudo orientalis*, a closely related leech species (20). The *Fusobacterium* OTU was previously reported in 16S clone libraries from the intestinum of *H. verbana* (14). The increase in Clostridial sequences observed in this study compared to previous studies is likely due to the improved DNA extraction techniques, which included a bead-beating step specifically optimized to increase clostridial cell lysis and therefore detection (34).

**Table 1.** Average percent of total 16S rRNA V4 sequences of core and common OTUs in ILF and intestinum samples from *Hirudo verbana* and *Macrobdella decora*

| Phylum | Genus | *Hirudo verbana* |  | *Macrobdella decora* |  | Accession No. |
| --- | --- | --- | --- | --- | --- | --- |
|  |  | ILF<sup>†</sup> | Intestinum | ILF | Intestinum |  |
| Bacteroidetes | Bacteroides | 1.37% | 0.38% | 39.82% | 40.48% | SAMN11823292 |
|  | Mucinivorans-like | 0% | 8.89% | 0% | 0% | SAMN11823282 |
|  | Millionella-like | 0.35% | 9.79% | 0% | 0.65% | SAMN11823299 |
|  | Mucinivorans | 58.20% | 16.83% | 0% | 0.00% | SAMN11823269 |
| Clostridia | Alkaliphilus-like | 0% | 1.11% | 20.42% | 28.62% | SAMN11823295 |
|  | Clostridium | 0% | 0% | 2.78% | 0.50% | SAMN11823294 |
|  | Papillibacter-like | 0% | 0% | 2.71% | 1.15% | SAMN11823296 |
|  | Butyricicoccus | 0% | 0% | 18.51% | 6.70% | SAMN11823293 |
| Alphaproteobacteria | Insolitispirillum-like | 0% | 21.02% | 0% | 1.67% | SAMN11823298 |
| Deltaproteobacteria | Desulfovibrio | 0.71% | 6.15% | 0.62% | 1.27% | SAMN11823279 |
| Gammaproteobacteria | Aeromonas | 18.07% | 15.88% | 8.10% | 9.15% | SAMN11823291 |

\* OTUs determined at the 97% confidence level. Those OTUs indicated as “-like” indicate the most closely-related genus listed in RDP.
<sup>†</sup> Core (present in ≥ 90% of samples) OTUs are highlighted in green. Common (present in ≥ 70% of samples) OTUs are highlighted in yellow. Core and common OTUS were determined for each species of leech and sampled organ individually.

In addition to the four most abundant taxa, common sequences (found in at least 70% of samples) in the *H. verbana* ILF included three Bacteroidetes, two *Bacteroides* (0 – 13%, median when present = 1.6%) and one *Millionella-like* OTU (0 – 2%, median when present = 0.4%); two *Proteocatella* (0 – 14.9%, median when present = 2.2%); and one *Desulfovibrio* (0 – 8.7%, median when present = 0.6%) (Table 1 and S2). Finding these additional OTUs in such a high percentage of animals is likely due to the much greater sequencing depth of Illumina technology as compared to those previously used. The additional *Bacteroides* and *Millionella-like* OTUs have not been described before in *H. verbana*. However, other researchers have previously noted that, although dominant, *Mucinivorans* may not be the only Bacteroidales associated with *Hirudo* leeches (14, 19, 35).

Sequences belonging to other taxa noted by previous researchers were also found in the 36 *H. verbana* at average concentrations below 1.5% of the total sequences from the ILF community: Firmicutes: *Erysipelothrix* (max = 5.1%, 17% of animals), *Vagococcus* (max = 1.2%, 17% of animals), and *Enterococcus* (max = 0.7%, 17% of animals); Alphaproteobacteria: *Ochrobactrum* (max = 0.4%, 6% of animals); Deltaproteobactera: *Desulfovibrio-like* (max = 5.9%, 39% of animals) and *Desulfovibrio* (0.1%, 3% of animals); Gammaproteobacteria: *Proteus* (max = 16.1%, 44% of animals) and *Morganella* (max = 5.2%, 53% of animals); Bacteroidetes: *Pedobacter* (max = 0.6%, 8% of animals) (14, 20, 21) (Table S2).

The low concentrations and intermittent presence of many of these previously identified bacteria highlights the importance of greater sequencing depth and a greater number of specimens tested. Our analysis of the microbial community in the ILF of *H. verbana* confirms that *Aeromonas* and *Mucinivorans* comprise the core microbiome with Clostridial species also dominant when present. Our analysis also reveals a more diverse rare *H. verbana* ILF microbiome than previously described as well as community members that are present in >70% of the animals.

### *M. decora* ILF Microbiome is Similar to *H. verbana* ILF Microbiome

One unique feature of the analysis of *H. verbana* is that these animals were obtained from leech farms that breed captive animals or maintain field-caught animals for months in captivity before shipping. Analyzing the microbiome of field-caught *M. decora,* would allow us to assess if the diversity of the microbiome could be greater in animals captured in the wild. The composition of the ILF microbiota from the 52 sampled *M. decora* was very similar to that of *H. verbana* in that it is dominated by *Bacteroides*, *Aeromonas*, and four Clostridiales spp (Table 1 and S2)*. Aeromonas* and *Bacteroides* sequences made up 16 – 73.5% (median = 51%) of the *M. decora* ILF microbiota while the Clostridiales spp accounted for another 22 – 98%. A *Bacteroides-like* OTU was present in *H. verbana* ILF at 0 – 2.1% of sequences (63% of animals) and sequences from only one of these Clostridiales species (*Alkaliphilus-like)* had been reported before from the *H. verbana* intestinum (14) (Table 1 and S2). To our knowledge, the remaining three Clostridiales species (*Papillibacter-like, Clostridium,* and *Butyricicoccus*) have never been identified from any site in *Hirudo* species.

Sequences from common OTUs (found in at least 70% of samples) in the *M. decora* ILF included one additional Bacteroidaceae and three Clostridiales (Table S2). These four OTUs made up an additional 0.6 – 12.8% of the ILF microbial community (median = 4.6%). The presence of more Clostridiales spp in the ILF microbiota is the primary cause for the observed increased alpha diversity in *M. decora* over *H. verbana* (Figure 2A). An increase in alpha diversity between *H. verbana* and *M. decora* ILF microbiota may have been a result of a ‘zoo effect’ in comparing farmed (*H.verbana*) to newly captured (*M. decora*) animals, an effect that has been well documented in mammals (36). This hypothesis will be addressed below when comparing the effect of feeding on the microbiota.

**Figure 2.**
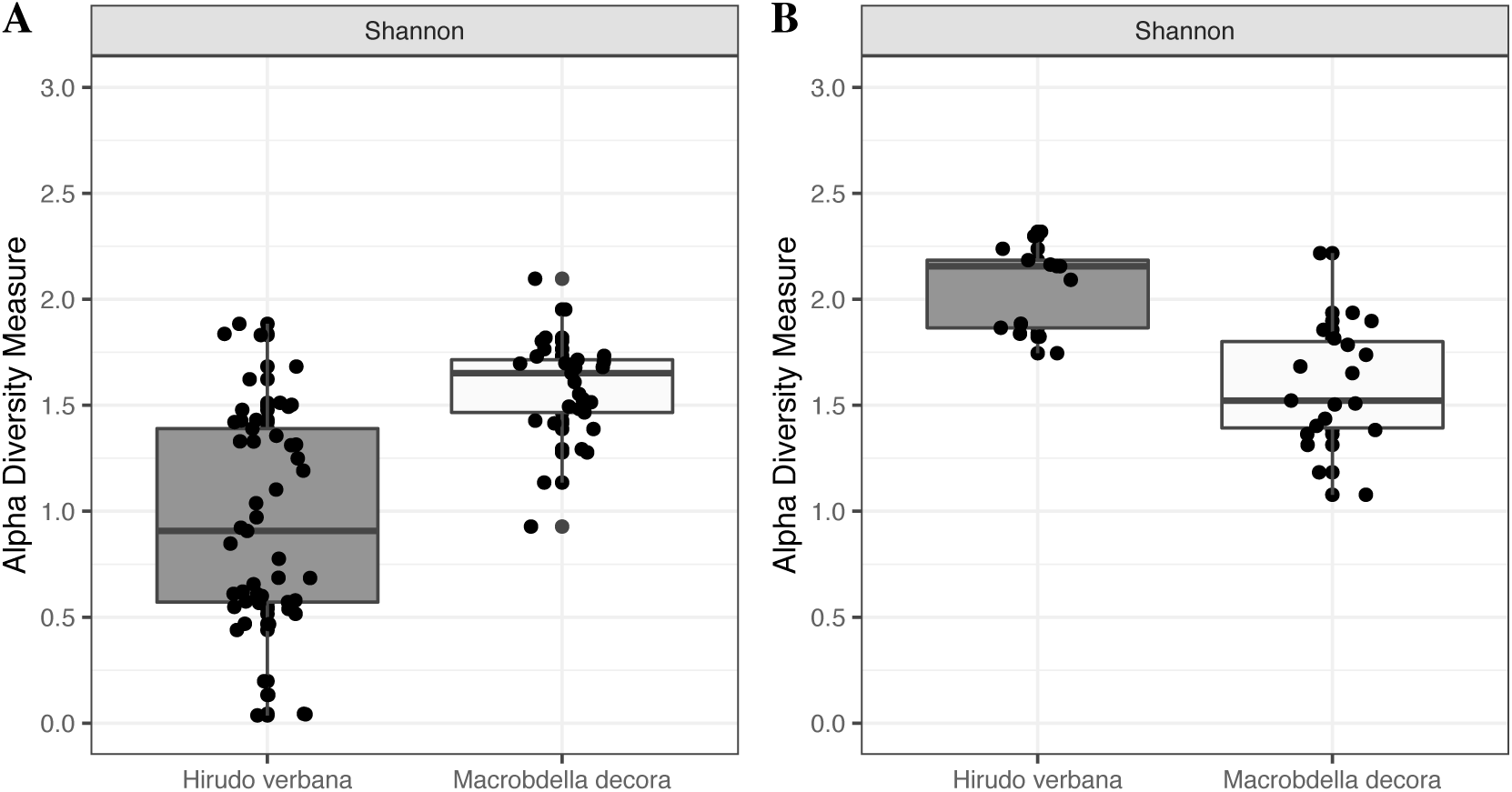
Shannon Alpha-diversity showing the microbiome of *Macrobdella decora* ILF is more diverse than that of *Hirudo verbana* but the opposite is true of the intestinum microbiomes. Calculations for leech A) ILF and B) intestinum samples. All samples were collected before feeding or greater than 28 days after feeding. Alpha diversity does not change between the ILF and intestinum of wild-caught *M. decora* (grey) but increases in farm-raised *H. verbana* (white).

### Microbiome of the Intestinum is Similar to that of the ILF

The intestinum microbiota from nine *H. verbana* samples was dominated by sequences from four OTUs: *Insolitispirillum-like* (0.2 – 34.6%, median = 25.4%), *Aeromonas* (2.3 – 24.7%, median = 16.6%), *Mucinivorans* (2 – 51.3%, median = 9.3%), and *Desulfovibrio* (1.7 – 15.3%, median = 4.1%) (Table 1 and S2). *Aeromonas* and *Mucinivorans* were also dominant members of the ILF community. Sequences from the *Insolitispirillum-like* and *Desulfovibrio* OTUs have been previously identified in the intestinum of *H. verbana* and *H. orientalis* (19, 20). The difference in microbial community between the *H. verbana* ILF and intestinum was marked by changes in the relative abundance of sequences from both dominant and common OTUs: *Insolitispirillum-like* (16.9 log_2_ fold change, p < 0.001), *Rikenella-like* (14.5 log_2_ fold change, p < 0.001), *Aquaspirillum-like* (13.8 log_2_ fold change, p < 0.001), *Mucinivorans* (−2.6 log_2_ fold change, p < 0.001) and *Proteocatella* (−11 log_2_ fold change, p < 0.001). The change in microbial composition between the ILF and intestinum may be directly related to the differences in organ functions between the two (erythrocyte storage vs meal digestion respectively (1)). The almost doubling of OTUs that were present in the intestinum versus those found in the ILF is consistent with previous reports (14, 19) of an increase in alpha diversity between the two organs in *Hirudo* spp (Figure 2B) and confirms that microbial colonization of the ILF is closely regulated.

In contrast, in *M. decora*, the alpha diversity between ILF and intestinum samples did not change significantly (Figure 2). The dominant OTUs of both ILF and intestinum from *M. decora* were *Aeromonas, Bacteroides*, *Butyricicoccus,* and *Alkaliphilus-like* (Table 1 and S2). In the intestinum (45 animals sampled), these OTUs accounted for 52.9 – 96.7% of the total sequences while in the ILF, these OTUs accounted for 73.2 – 98.8% of the sequences (Table 1 and S2). Sequences from common OTUs (found in at least 70% of samples) in the *M. decora* intestinum included additional *Bacteroides*, *Proteocatella-like, Desulfovibrio,* and two *Alkaliphilus-like* OTUs. An explanation for the observation that the microbiome of *M. decora* ILF is not as reduced as that observed in *H. verbana* may a result of maintaining or raising the animals in a farm.

The dominant *Alkaliphilus-*like OTU from *M. decora* was previously found in the *H. verbana* intestinum (14) but was detected only in three intestinum and three bladder samples from *H. verbana* processed in this study. All of the other OTUs common in the *M. decora* intestinum were not found in *H. verbana* samples. In the intestinum, like in the ILF, OTUs defined to the genus level are not the same but when identified to the order level community composition is conserved.

### *H. verbana* Bladder Community Contains Three Core OTUs

16S rRNA V4 deep sequencing of bladders from ten *H. verbana* animals revealed a more complex community than had been previously reported through RFLP and 16S rRNA gene sequencing (30). Kikuchi et al reported the *H. verbana* bladder contains a stratified community including Alphaproteobacteria *(Ochrobactrum),* Betaproteobacteria *(Comamonas-like & Sterolibacterium-like),* Deltaproteobacteria *(Bdellovibrio),* and Bacteroidetes *(Niabella & Sphingobacterium)* with *Ochrobactrum, Comamonas-like, Bdellovibrio, Niabella,* and *Sphingobacterium* found in > 90% of animals tested (30). The deep sequencing carried out in the current study on bladders from ten animals expanded these results by identifying Alphaproteobacteria *(Ochrobactrum, Aminobacter, Ensifer, Insolitispirillum-like, Phreatobacter-like),* Betaproteobacteria *(Ramlibacter, Pelomonas, Variovorax, Acidovorax, Ralstonia),* Deltaproteobacteria *(Bdellovibrio-like* and *Desulfovibrio),* Bacteroidetes *(Niabella, Pedobacter,* and *Flavobacterium),* and *Spirochaetes (Spirochaeta-like)* in *H. verbana* bladders although only *Ochrobactrum, Ramlibacter* (previously *Comamonas-like*)*, Bdellovibrio-like,* and *Pedobacter* were recovered frequently enough (in ≥ 90% of samples) to be considered part of the core (Table 2).

**Table 2.** Presence of bladder OTUs in total 16S rRNA V4 sequences from *Macrobdella decora* and *Hirudo verbana* bladder samples.

| Class | Genus | GenBank<br>Accession No. | *Hirudo verbana* |  | *Macrobdella decora* |  |
| --- | --- | --- | --- | --- | --- | --- |
|  |  |  | Median<sup>‡</sup> | Max<sup>§</sup> | Median | Max |
| Alphaproteobacteria | Ochrobactrum | SAMN11823287 | 15.46% | 94.99% | 0% | 12% |
|  | Ensifer | --- | 0.18% | 13.18% | 0.37% | 13.14% |
|  | Aminobacter | --- | 0.40% | 2.92% | N.D. | N.D. |
|  | Azospirillum | --- | N.D. | N.D. | 15.13% | 28.86% |
|  | Azospirillum | SAMN11823283 | N.D. | N.D. | 19.08% | 45.92% |
|  | Phreatobacter-like | SAMN11823288 | 0.86% | 9.52% | 3.23% | 21.54% |
| Betaproteobacteria | Methylopumilus-like | SAMN11823285 | N.D. | N.D. | 6.68% | 17.40% |
|  | Ramlibacter | SAMN11823289 | 20.81% | 75.62% | 20.94% | 38.93% |
| Deltaproteobacteria | Bdellovibrio-like | SAMN11823266 | 2.22% | 20.09% | N.D. | N.D. |
| Bacteroidetes | Niabella | SAMN11823286 | 0.63% | 9.20% | N.D. | N.D. |
|  | Pedobacter | SAMN11823267 | 18.48% | 44.24% | N.D. | N.D. |

<sup>‡</sup> Median percentage of total 16S V4 rRNA sequences, when OTU is present. Core (present in $\geq 90\%$ of samples) OTUs are highlighted in green.
<sup>§</sup> Maximum percentage of total 16S V4 rRNA sequences.

The increase in OTUs identified in the *H. verbana* bladder was likely due to sequencing three times as many animals as in the previous study as well as sequencing to a much greater depth. The expanded community contains sequences from an additional four Alphaproteobacteria, three Betaproteobacteria, one Deltaproteobacteria, and one Spirochaete. Comparing the 16S rRNA V4 sequences to the 16S rRNA sequences produced by Kikuchi et al, indicates that the *Ochrobactrum, Ramlibacter*, *Niabella,* and *Pedobacter* (previously *Sphingobacterium*) OTUs are the same between the two studies (Table S2). However, the previously-identified *Sterolibacterium-like* and *Bdellovibrio* OTUs did not have any similar sequences in the new dataset. The increased number of identified OTUs in conjunction with the small number of OTUs identified as core suggests that there is large variation in the minor members of the bladder communities between individual animals. Further research would be required to determine if this large variation is also apparent between bladders of an individual animal.

### *M. decora* Bladder Community

The 16S rRNA V4 sequencing of *M. decora* bladder from 20 animals identified sequences belonging to *Alphaproteobacteria (Ochrobactrum, Ensifer, Rhizobium, Rhizobium-like, Azospirillum, Phreatobacter-like, Sphingomonas, Rhodopseudomonas),* Betaproteobacteria *(Ramlibacter, Methylopumilus-like, Bacteriovorax-like, Pandoraea,* and a Rhodocyclaeae sp), and *Deltaproteobacteria (Bdellovibrio, Cystobacter-like, Desulfovibrio,* and *Bacteriovorax-like).* Of these identified taxa, only *Ramlibacter, Methylopumilus, Phreatobacter-like,* and *Azospirillum* were considered core (Table 2). The *Ramlibacter, Ochrobactrum,* and *Sphingobacterium* OTUs sequenced from *M. decora* are the same as those identified from *H. verbana* bladders (30) (Table S2) and suggest a conservation of bladder symbionts at the genus level (97% sequence similarity) despite geographic and evolutionary separation.

Surprisingly, *Aeromona*s, *Bacteroides,* and Clostridial species were also detected in the bladder sequences from *M. decora* and *Aeromonas*, *Mucinivorans,* and Clostridial species were identified in the bladder sequences from *H. verbana*. Two possibilities seem likely to be responsible for this observation: (i) ILF symbionts are also found in the bladder and nephridia or (ii) contamination from the ILF occurred during dissection, PCR, or sequencing (37). The contamination seemed especially likely in animals that had been recently fed as the filled crop is easily punctured during dissection. In addition, previous data from *H. verbana* bladder communities suggested that it was unlikely that these two dominant ILF taxa were present in the bladder (30).

In an effort to confirm the 16S V4 rRNA deep sequencing results, FISH was used to detect specific taxa in the bladder. Unexpectedly, low levels of Aer66-binding cells were detected in cells lining the *M. decora* bladder/nephridia of animals 4 and 7 days after feeding (Figure 3). While this observation confirms the deep sequencing results, it raises questions to the origin of these *Aeromonas* cells. A SILVA probeCheck (38) of the Aer66 probe confirms that it should be specific for *Aeromonadaceae*, interestingly, there is only a single base pair mismatch to the *Ochrobactrum* sequence. Thus, it remains unclear if this probe is binding to the highly abundant *Ochrobactrum* or *Aeromonas*. Because the Bacteroidetes probe used in this study also targets *Pedobacter* and *Flavobacterium* and may cross-react with *Phreatobacter-like*, we cannot conclusively determine whether the Bacteroidetes symbiont exists within the *M. decora* bladder.

**Figure 3.**
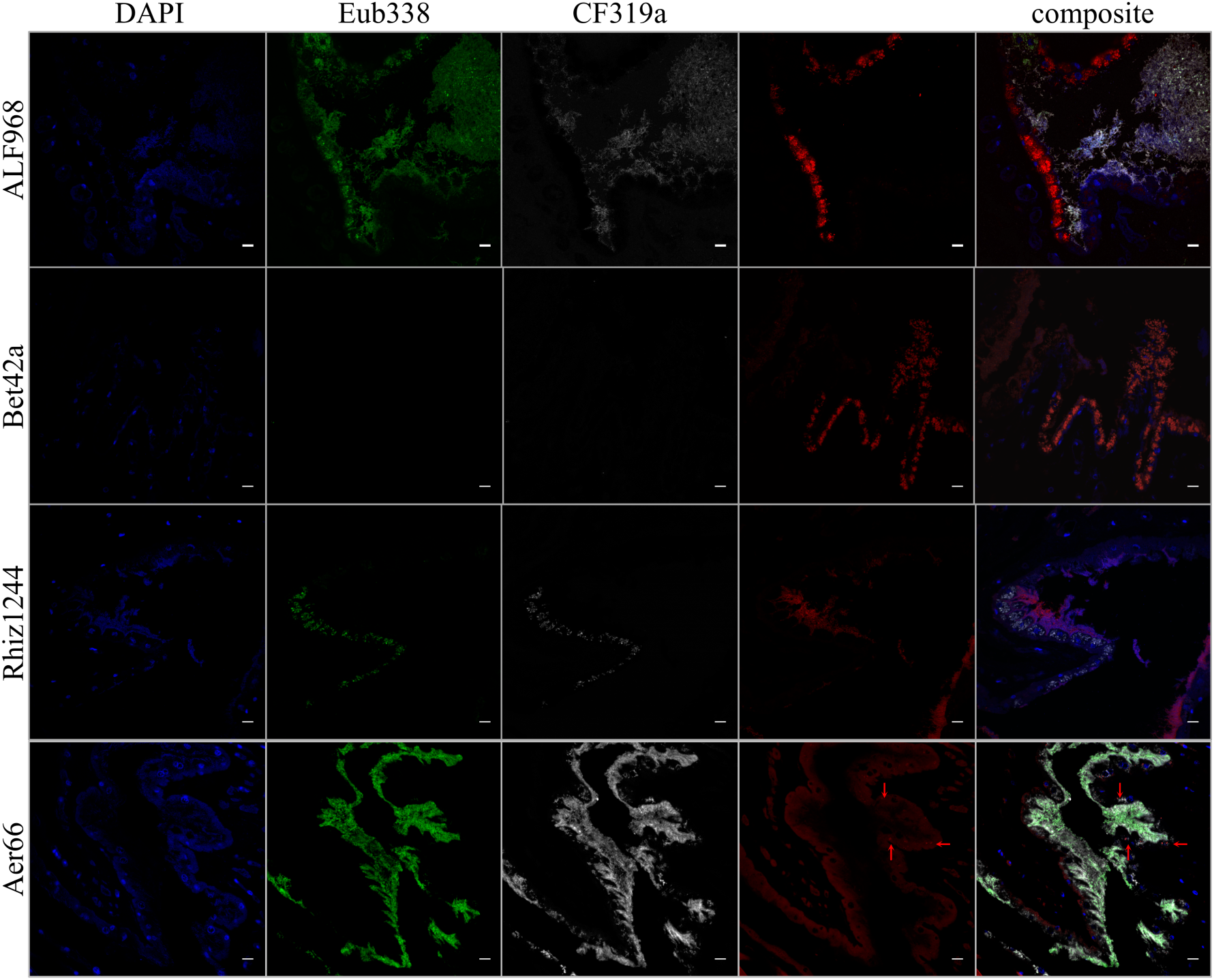
False-color FISH micrographs of *Macrobdella decora* bladder samples. From left to right, columns contain images with probes DAPI (blue – Eukaryotic DNA), Eub338 (green – Eubacteria), CF319a (white: Bacteroidetes *– Mucinivorans, Bacteroides, Niabella, Pedobacter*), (red), and composite. From top to bottom, the fourth (red) column contains images with probes Alf968 (Alphaproteobacteria *– Ochrobactrum, Azospirillum, Phreatobacter-like*), Bet42a (Betaproteobacteria – *Methylopumilus-like, Ramlibacter*), Rhiz1244 (Rhizobiales – *Ochrobactrum, Phreatobacter-like*), Aer66 *(Aeromonas)*. The *M. decora* bladder is colonized in a stratified manner with intracellular *Azospirillum*, epithelial-associated Betaproteobacteria, and *Phreatobacter-like* and *Niabella* in the matrix. *Aeromonas* (red arrows) is present associated with eukaryotic cells, suggesting that they have been carried here by leech hemocytes and are not normal flora for the bladder. Bars = 10*μ*m.

### Physically Stratified Bacterial Community in *M. decora* Bladders

FISH imaging of *M. decora* bladders suggests that most Alphaproteobacteria (*Ochrobactrum* and *Phreatobacter-like*) occur intracellularly in the epithelial cells lining the interior of the bladder, while Rhizobiaceae (*Aminobacter* and *Ensifer*) occur in the matrix of the bladder contents. Betaproteobacteria (*Methylopumilus-like* and *Ramlibacter*) occur in close association with the *M. decora* bladder epithelial cells (Figure 3). This layered community is consistent with localizations reported previously in *H. verbana* bladders (30). Together, this data confirms that the *M. decora* bladder community composition and distribution are similar to those of *H. verbana* despite differences in specific OTUs and suggests an evolutionary conservation between these hosts and their symbionts.

### Seasonal Changes Affect the Wild ILF Microbiome

The *M. decora* animals used in this study allowed us to evaluate how a wild leech’s microbiome changes during the year. Animals collected during warm months (June-September) had similar ILF and intestinum microbiomes. However, animals collected in October (the beginning of cold months) contained significantly different ILF microbiomes from those collected in April (end of cold months) or during warm months (p = 0.003) (Figure 4A). Those collected in April showed the least variation between individuals.

**Figure 4.**
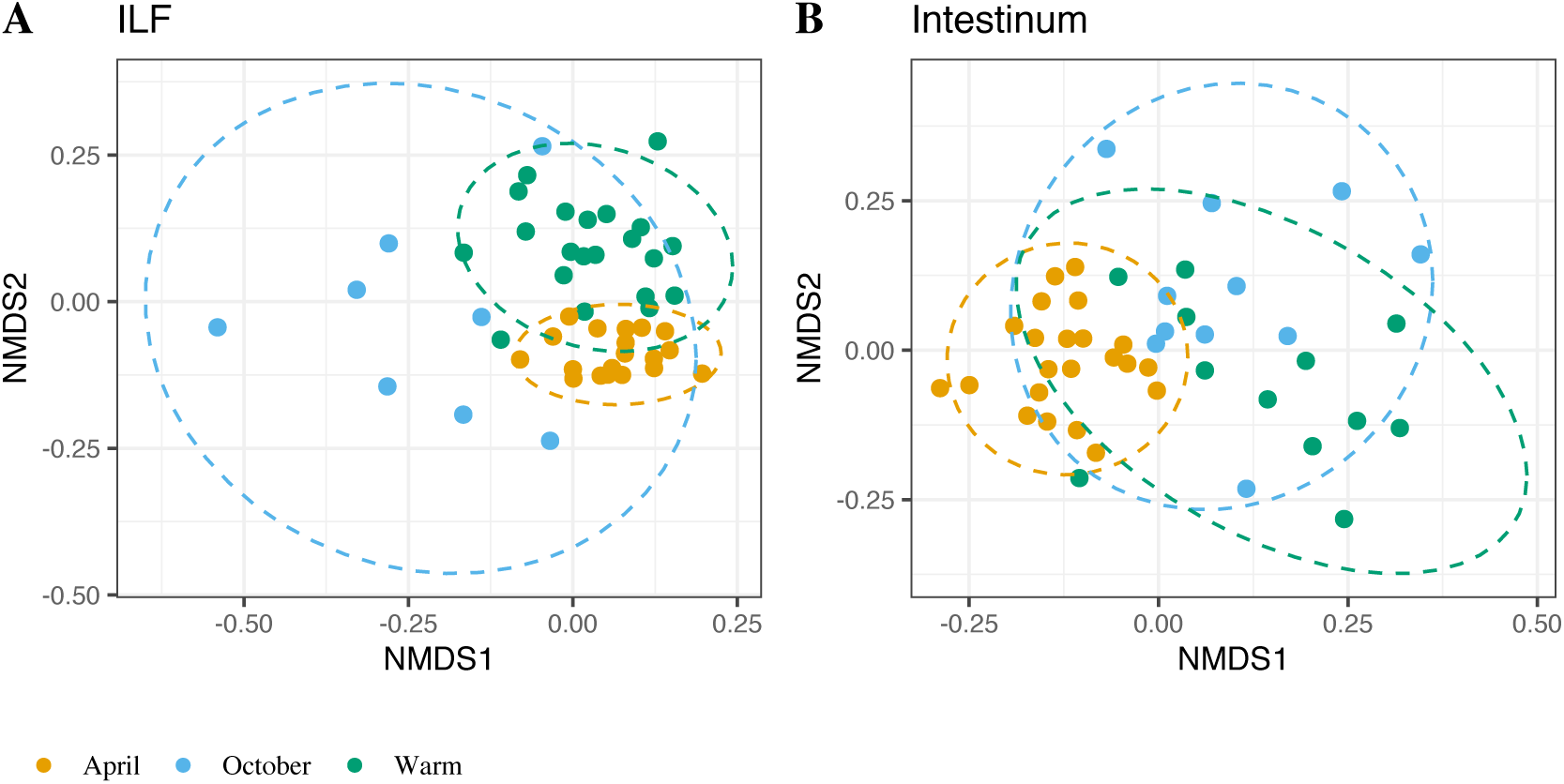
Unifrac-calculated NMDS plot of wild-caught *Macrobdella decora -* associated microbiota shows that month of collection affects gut microbiome. Animals were collected in three seasons: late cold (April - light blue), warm (June - pink, July - red, August - brown, and September - brown), and early cold (October - dark blue). A) ILF samples. ILF microbiota from animals collected in October were significantly different from that of animals collected in April or warm months. B) Intestinum samples. Intestinum microbiota from animals collected in April were significantly different from that of animals collected in warm months or October. Ellipses drawn at 95% confidence interval.

Among intestinum samples, those from animals collected in April were significantly different from those collected in October or during warm months (p = 0.003 and p = 0.009 respectively) (Figure 4B). However, the difference in intestinum microbiome between animals collected in warm months were not significantly different from those collected in October (p = 0.123). The seasonal difference in the microbiome composition was detected over multiple years with animals collected during warm months being more similar to those collected in warm months of other years than to those from animals collected in April or October of the same year.

### Feeding Affects the ILF Bacterial Community

Sanguivorous leeches are opportunistic ectoparasites that consume several times of their body weight in a single feeding and rest for months between feeding events (7). Following a feeding event, the abundance of *Mucinivorans* and *Aeromonas* in the *H. verbana* ILF increases (13, 15)). Monitoring of bacterial communities after a laboratory-administered, sterile sheep blood meal in *M. decora* revealed a significant decrease in alpha diversity in ILF by 2 days after feeding (DaF) that rebounded by 30-90 DaF (Figure 6). At 2 and 4 DaF the communities in the ILF were significantly different from those at any other time after feeding (p = 0.009 and p = 0.003 respectively). A similar change in community was observed in *H. verbana* where *Mucinivorans* and *Aeromonas* populations increased after feeding (15). Interestingly, *M. decora* seemed to be divided into two groups, those that recovered quickly (by 4 – 7 DaF) and those that recovered more slowly (> 7 DaF). These groups can be seen in the dichotomy in the violin plot of Bray-Curtis distances presented in Figure 6B.

By 30 DaF, the alpha diversity and common members of the ILF communities of both *H. verbana* and *M. decora* returned to levels indistinguishable from those in unfed animals (Figure 5 and 6). This would suggest that a single blood meal and maintenance in the artificial lab environment was not sufficient to significantly change the composition of the gut community. However, it should be noted that animals maintained for longer than 90 days before sampling did appear to have a reduced variation between individuals (data not shown). This would suggest that a zoo effect is minimal, although a slight decrease in biodiversity from reduction in the rare microbiome members should be anticipated, as has been described in mammals (39). The abundance of sequences from dominant members of the gut microbial community overall appeared to return to levels comparable to those of unfed animals within 30 DaF (Figure 5).

**Figure 5.**
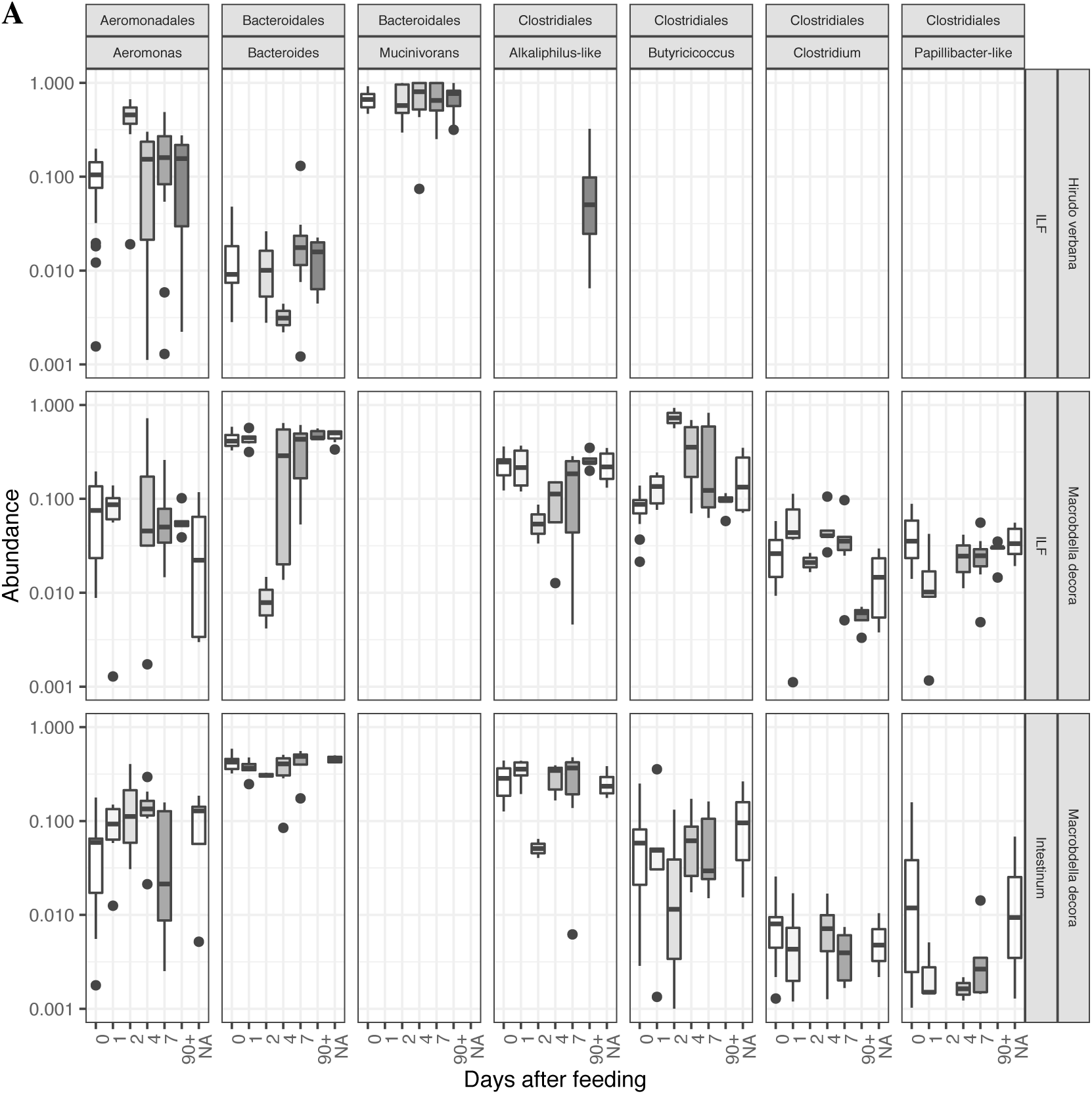
Box and Whisker plot of leech-associated gut microbiota showing that the prevalence of taxa in the ILF appears to be greatly affected within ∼48h of a blood meal, while the prevalence of taxa in the intestinum maintains relative stability. The prevalence of 9 core gut taxa from deep-sequenced microbiomes of A) *Hirudo verbana* ILF, B) *Macrobdella decora* ILF, and C) *M. decora* intestinum were assessed at 7 time points after a blood meal (0, 1, 2, 4, 7, 30, and 90+ days after feeding). See Supplemental Table 1 for number of samples analyzed at each time point.

**Figure 6.**
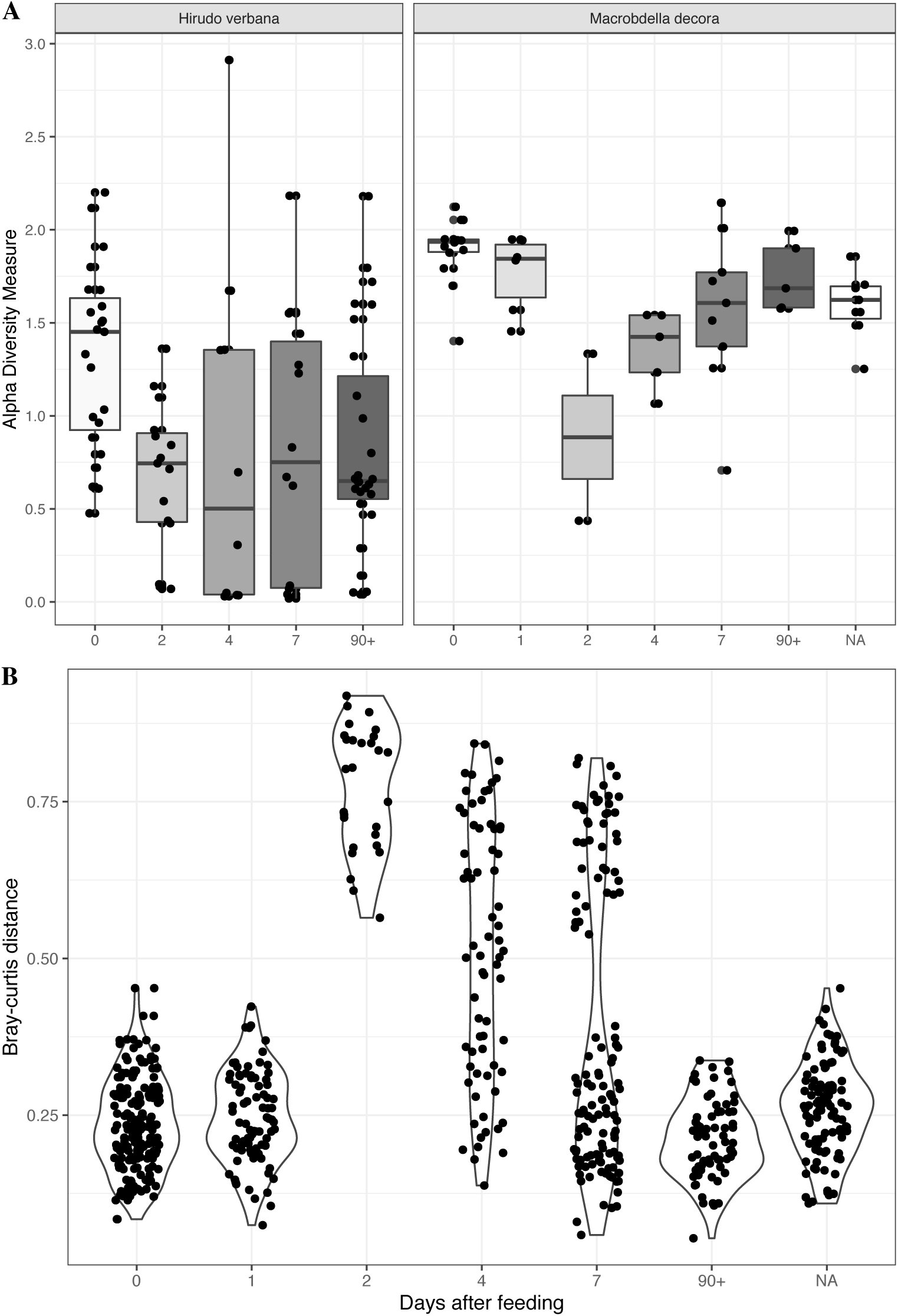
*Macrobdella decora* ILF diversity changes with time after feeding. A) Shannon diversity index calculates a drop in ILF microbiome diversity by 2 days after feeding (DaF) that rebounds by 7 – 30 DaF. B) Bray-curtis distance calculation shows that ILF microbiomes at 2 – 7 DaF are significantly different from those of unfed animals (0 DaF) and that the community rebounds at 30 – 90 DaF. Note the appearance of two sub-populations of ILF samples that i) appears to rebound by ∼4 DaF and ii) appears to require > 7 DaF to rebound.

The dominant *Bacteroides* and *Aeromonas* symbionts identified by deep sequencing were additionally observed through FISH imaging. Immediately after capture, the microbial population in the *M. decora* crop was below the limit of detection for FISH, although rare *Aeromonas* cells were occasionally found (Figure 7). After feeding a blood meal the microbial population increased at 4 and 7 DaF (Figure 7) with Bacteroidetes forming large microcolonies by 4 DaF and *Aeromonas* pelagically spread throughout the crop by 7 DaF. A similar pattern was previously observed and quantified in *H. verbana* (15). Interestingly, some *Aeromonas* at 7 DaF were found associated with eukaryotic cells within the crop. These cells are likely circulating immune cells from the leech host. Phagocytosis of *Aeromonas* by leech hemocytes was previously observed in *H. verbana* when the *Aeromonas* strain lacked a functioning T3SS but not for the wild type (40).

**Figure 7.**
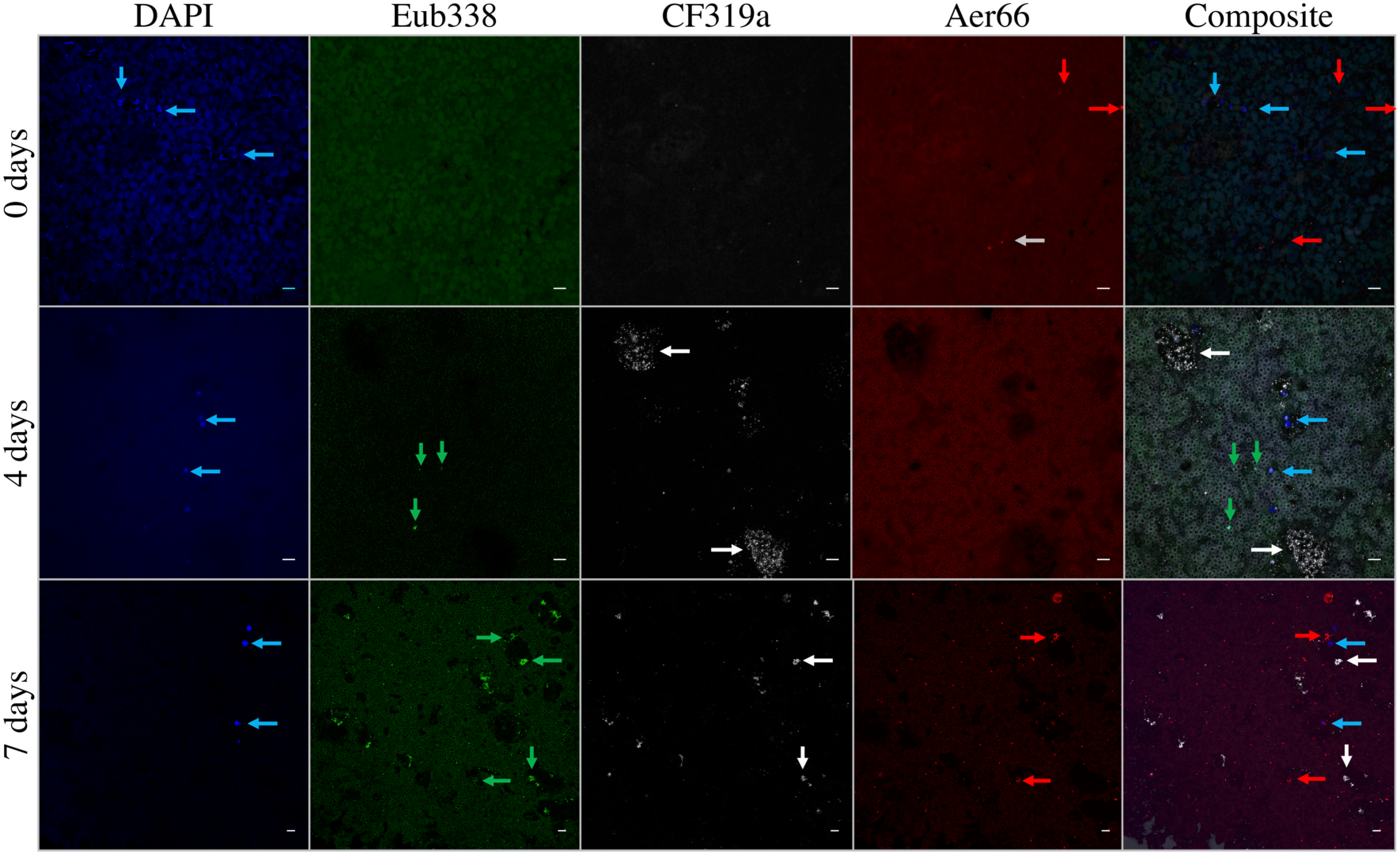
False-color FISH micrograph of *Macrobdella decora* ILF. From left to right, columns contain images with probes DAPI (blue – Eukaryotic DNA), Eub338 (green – Eubacteria), CF319a (white: Bacteroidetes *– Mucinivorans, Bacteroides*), Aer66 (red – *Aeromonas*), and composite. From top to bottom, the rows contain images from animals sacrificed 0 days (wild-caught animals), 4 days, and 7 days after a laboratory-administed sterile blood meal. Blue arrows indicate eukaryotic cells (most likely leech hemocytes), green arrows indicate notable bacteria, white arrows indicate Bacteroidetes microcolonies, and red arrows indicate *Aeromonas.* Background fluorescence is from crop contents, especially the blood meal at 4 and 7 days. Few bacteria are present in the crop of wild-caught, unfed animals. Bacteroidetes colony expansion occurs by 4 days after feeding, while *Aeromonas* prevalence increases by 7 days after feeding. Other bacteria are present at 4 days after feeding, however their numbers appear to be overwhelmed by Bacteroidetes and *Aeromonas* growth at 7 days after feeding. In animals 7 days after feeding, *Aeromonas* occasionally are found associated with hemocytes. Bars = 10*μ*m.

### Feeding Does not Affect the Intestinum Bacterial Community

Unlike the bacterial community present in the ILF, the intestinum bacterial community appears to be much more stable. No significant change between any of the days after feeding were observed (p ≥ 0.15). This suggests that growth of any single member of the community is matched by other members. More likely, the intestinum may be a more constant environment with a regular inflow and outflow that is not as much affected by sporadic feeding events. It is also possible that the intestinum serves as a reservoir from which new blood meals may be seeded with symbionts after ingestion.

### Geographic Location

Because *M. decora* were collected from multiple ponds in nature, we were able to compare the microbial communities of animals from the same species but distant geographic locations. There was no significant difference between microbial communities in the ILF of *M. decora* from CT, MA, NY, and VT (p ≥ 0.222) (Figure s1). Statistical analyses on geographic effects were performed only on samples from MA and CT due to the lower sample number for samples from NY and VT.

The microbiome from the intestinum of MA animals was significantly different from that of CT animals (p = 0.006), but interestingly the ILF microbiome was indistinguishable between the two leech populations (Figure s1). The difference in microbial communities between the ILF and intestinum was significant in animals from MA (p = 0.006), but not in animals from CT (p = 0.084), and was associated with a decrease in *Alkaliphilus-like* (p = 0.022) and increase in *Papillibacter-like* (p = 0.001) OTUs. In the ILF, these OTUs make up ∼ 8.2% of the total ILF sequences while in the intestinum they make up less than 1.6%. This difference between ILF and intestinum microbiota is almost imperceptible in animals from CT. The slight differences in Clostridiales species between MA and CT animals suggests a site-specific variation in less abundant community members within *M. decora*, however the site-specific dominant symbionts and core community remained the same and indicate a conservation of symbionts despite geographic separation.

## DISCUSSION

In this study we have used deep sequencing of the V4 region of the 16S rRNA gene and FISH to characterize the microbiome from three organs in the North American medicinal leech, *Macrobdella decora*. At the taxonomic level of order, the microbiomes described from wild-caught *M. decora* appeared very similar to those from *Hirudo verbana*. However, at the genus level, the microbiomes were easily differentiated between the two host leech species. This is similar to findings in other animal models where often the host-specific genera show an evolutionary pattern that mimics that of the host’s evolution (41-45).

In both leech species, the gut microbial communities are dominated by *Aeromonas,* Bacteroidales, and Clostridiales species. This is consistent with previous findings that Bacteroidetes and *Aeromonas* are common but not obligate in the gut of animals and insects fed on blood (46-51). Siddall et al. have shown that many sanguivorous leeches maintain a gut symbiosis with *Aeromonas* and Bacteroidetes species (8). Because of the difficulty of distinguishing *Aeromonas* species based solely on 16S sequencing data (52, 53), we are unable to determine whether the *Aeromonas* species from *M. decora* is the same as that from *H. verbana*.

In-depth genetic analyses of an *Aeromonas veronii* strain isolated from *H. verbana* has shown that the ability to grow in blood is not sufficient for leech gut colonization, but rather that a number of colonization factors such as secretion systems (54), carbon starvation response (55), oxidative stress response inhibition (7), and heme acquisition processes (56) are critical in enhancing leech gut colonization. New studies comparing genomic variation between *Aeromonas* strains isolated from sanguivorous leech species to those of non host-associated environmental strains may be used to further identify genes critical for host colonization and persistence.

In addition to confirming a similar core and common microbiome between the two leech hosts, our study also identified a rich rare microbiome. Previous research in a variety of environments has concluded that although a relatively small number of OTUs may dominate samples, low-abundance populations may be responsible for driving changes in observed phylogenetic diversity (57, 58). Rare species may have a significant role in metabolic cycles and may be a hidden driver of ecosystem functioning (59). Because dominant and core microbes are easier to identify and predict, rare microbes may be overlooked keystone species responsible for regulating the community’s development and function (59). In our study, the ability to examine an increased number of animals, to examine three body sites from each animal, and to sequence to a much greater depth resulted in the detection of an increased level of inter-animal variations in the less abundant microbiota and suggest a greater diversity in the rare microbiome than previously predicted.

We were also able to observe the effect of geographic isolation on the less abundant members of the gut microbial community in *M. decora*. The very slight but statistically significant differences in community composition between animals from MA and CT indicated that the relationship with dominant symbionts is consistently conserved despite geographic separation of ∼100 km. It is possible that this is due to a small sample size as we compared 13 unfed *M. decora* from MA and seven from CT and previous studies of synthetic and natural microbial communities found that insufficient sampling could yield an artificial difference between geographically separated communities (60, 61). As has been noted previously in the termite gut (58), the difference in community composition is based on minor members of the community and is reminiscent of a similar pattern observed in community differences between suppliers of *H. verbana* (data not shown). We therefore hypothesize that the conserved members of the community have been evolutionarily maintained in hirudinid leeches despite species-diversification and continent-dividing events.

Not only was the core gut microbiome similar between *M. decora* and *H. verbana*, but also the dynamic response of these symbionts in response to feeding a blood meal. FISH imaging confirmed similar colonization patterns of the ILF of *Aeromonas* and Bacteroidetes from *H. verbana* and *M. decora*. Although part of the decrease in alpha diversity in the leech ILF microbiome post blood meal consumption may be due to innate immune properties of the blood meal (62), in a number of vertebrate species it has also been shown that extreme feeding or fasting events do result in decreased alpha diversity (63, 64). In *H. verbana*, it has been suggested that the *Aeromonas* symbiont may provide additional antimicrobial peptides responsible for restricting bacterial colonization in addition to host-produced antimicrobial peptides (65). Because the core community and post-feeding community dynamics are similar in *M. decora*, it is reasonable to assume that a similar process may also occur in this host species.

While our laboratory-maintained animals were tracked for 30-90 days after feeding, in the wild leeches are thought to go for even longer periods of time between feedings as the animals are inactive during the cold months. This extended period between feedings likely resulted in a starved phenotype in animals collected in April. Our results are consistent with the long starvation period occurring during the cool months affecting the gut microbial community of *M. decora* and the relative nimiety of the warm months where feedings may occur regularly. The observed seasonal effect may also be driven by the sequencing of transient environmental microbes from within the leech crop, as has been observed in a recent limited survey of wild *H. verbana* (17).

Changes in gut microbial populations as a result of irregular feeding events has been observed in other animals too. Other researchers have observed Clostridial populations increase rapidly in response to feeding while Bacteroidales numbers increase at a slower rate and high levels of Bacteroidales or Rikenellaceae are indicative of long periods between feedings in avian, reptilian, and mammalian hosts (63, 64, 66). In our experiments, a higher numbers of Bacteroidales sequences were found in the ILF of *M. decora* collected in April when compared to those collected in June-October. The proportion of Bacteroidales sequences additionally decreased in leeches immediately after feeding (when the number of Clostridiales sequences increased). FISH imaging did indicate that Bacteroidales increase in number four days after feeding, suggesting that the change in sequence ratio is due to a combination of rapid replication of Clostridiales and a much slower replication of Bacteroidales. This consistency with community dynamics observed in other animal models then suggests that leeches may be good models to predict microbial changes due to intermittent feedings or extended periods of starvation.

In addition to the conservation of *Aeromonas*, Bacteroidetes, and Clostridial gut symbionts, the bladder taxa *Comamonadaceae, Ochrobactrum,* and *Nubsella* appear to have also been evolutionarily maintained within Hirudiniform leech bladders. They are common in annelid bladders as are *Bordetella, Methylophilus, Achromobacter, Variovorax, Azospirillum, Mesorhizobium, Phyllobacterium, Rhizobium, Desulfovibrio, Pedobacter, Spirochaeta* (28). The taxa from the bladders of both *H. verbana* and *M. decora* were very similar to those found in other annelids, which include Alphaproteobacteria *(Ochrobactrum, Azospirillum,* Brucellaceae, Rhizobiaceae, Rhodospirillaceae, & Bradyrhizobiaceae), Betaproteobacteria (Comamonadaceae, Methylophilaceae, & Rhodocyclaceae) and Bacteroidetes *(Pedobacter)* (28, 29). This observation suggests an even greater conservation than that observed in digestive symbionts as the conservation has occurred at the genus level in medicinal leeches and at the family level over multiple evolutionary events in the annelids.

The *Ramlibacter* (previously *Comamonas-like*)*, Ochrobactrum,* and *Pedobacter* (previously Sphingobacterium) OTUs sequenced from *M. decora* are the same as those identified from *H. verbana* bladders (30). However, the *Pedobacter* sequenced in this study does not seem to be the same that Ott isolated from the mucus castings of adult *H. verbana* (67). This may indicate a difference between the bacteria on the outside of the leech and those adapted to colonize the bladder of the leech, as has been observed in the specialization of *Aeromonas* strains inside the leech gut (27, 54-56, 68). Further research is required to determine how these two leech species are able to maintain such specific bladder symbioses and by what means the symbionts are efficiently transmitted between animals.

Based on the deep-sequencing absence of *Niabella, Pedobacter, Ramlibacter*, *Ensifer, Phreatobacter,* and *Aminobacter* sequences in ILF and intestinum samples, the low abundance of *Ochrobactrum* sequences in ILF samples (max = 0.4%), and the absence of any ALF968-, Rhiz1244-, or Bet42a-binding cells in FISH images (data not shown) we hypothesize that these genera are specific to the bladder. Presence of these OTUs in an ILF sample might be due to contamination during dissection or a diseased state of the animal. This has occurred in previous studies where bladder-specific symbionts were found in the ILF and intestinum of 3 wild *H. verbana* (17) and *Pedobacter* and *Ochrobactrum* were cultured from intestinum samples (14, 69). The fact that, in the Worthen study, *Pedobacter* and *Ochrobactrum* clones were only recovered from recently-fed animals supports the possibility of sample contamination during dissection (recently fed animals have larger bladders). Because of the friability of leech tissue and the examples of bladder symbionts appearing in ILF and intestinum samples, careful dissection technique is critical to reduce contamination during sample acquisition.

Our research has identified a conservation of gut and bladder symbioses in medicinal leech species from Europe and North America. While the community structure of *H. verbana* and *M. decora* microbiomes have specific similarities, the differences that exist between the two can be exploited in the future to gain a better understanding of host/symbiont co-evolution. Not only are the gut symbionts conserved, but their dynamics in response to a blood feeding are similar as well. From our data we can hypothesize that the gut microbiome composition is affected by periods of starvation and torpidity. In *H.verbana*, the simple gut community helps to digest the blood meal (7) and protect the host from invading bacteria (65). It is likely that the microbial communities in the gut of *M. decora* perform very similar functions. Future work should focus on isolating dominant members of both symbioses to compare conserved metabolic and colonization abilities and to test for host species specialization. Through understanding molecular and mechanistic studies enabled by simple systems, especially in evolutionarily conserved biological processes and metabolic capabilities, one can form predictions that are more challenging to test in more complex communities.

## METHODS

### Animals

*M. decora* were collected from ponds located in Storrs, CT (41’49’3.074”N, 72’15’32.704”W), Groton, MA (42’35’26.993”N, 71’32’25.63”W), Caroga, NY (43’11’28.684”N, 74’28’10.026”W), and Mount Snow, VT (42’58’25.3”N, 72’55’47.9”W). If sacrificed within 1 week of collection, animals were maintained in native pond water.

*H. verbana* were purchased from Leeches U.S.A. (Westbury, NY U.S.A.) and BBEZ (Biebertal, Germany) and maintained as described below.

### Husbandry

Leeches were maintained in circular tanks (up to 10 animals/tank) in an environmental chamber with 12/12h day/night cycles at 25/23’C respectively. Sterile dilute instant ocean (DIO) consisting of 34 mg/L instant ocean salts (Aquarium Systems, Inc Mentor, OH U.S.A.) in nanopure water was changed weekly or when the animals vomited or an animal in the tank died. Tanks contained a minimum of one autoclaved rock.

Tanks were cleaned by completely emptying the old water and performing multiple complete, small volume, water changes until water remained clear for at least 20 min after the last change. Tanks were then filled to previous level and returned to environmental chamber.

### Feeding

Animals were fed as described previously (4). Sheep blood with heparin anticoagulant was purchased from Lampire Biological Laboratories (Pipersville, PA U.S.A.). Animals were fed in groups of 1-3 animals on sterile 50 mL Falcon tubes containing 30 mL sheep blood, warmed to 37’C, and covered with Parafilm (American National Can, Greenwich, CT U.S.A.). All animals were allowed to feed to satiety and then let sit in sterile DIO for ∼30 min before handling after feeding.

### Dissection

Animals were removed from the tank and the anterior end was tied with string to inhibit regurgitation before narcotizing in 70% ethanol (13). Animal was then treated with RNase away and rinsed with molecular biology grade water. An additional string was tied around to bissect the animal into anterior and posterior fractions.

Bladder: A small primary incision was made immediately lateral to the dorsal lateral dextral stripe and immediately posterior to the second string. Bladders were identified and carefully dissected so as to minimize contamination from ILF. Bladders were briefly rinsed in sterile phosphate buffered saline, PBS, then placed in sterile bead-beating tube. Each sample consisted of 1-3 bladders from the same leech. In this manuscript, the term ‘bladders’ includes bladders and the attached nephridia.

ILF: To access the crop contents, a secondary central incision was made immediately below the second string. Approximately 100 *μ*L ILF was collected as it was released and placed in a sterile bead-beating tube. In animals with especially low volumes of ILF, a sterile pipette tip was used to gently scrape the inside of the crop.

Intestinum: To access the intestinum, a second incision was made immediately lateral to the anus. The intestinum was carefully dissected from the anus and consisted of ≥1cm intestinum with contents. Samples were placed in bead-beating tubes or 1.5 mL microcentrifuge tubes and snap frozen in liquid nitrogen before storage at −80’C.

### Tissue Fixation and Embedding

After bladder, ILF, and intestinum sample were collected a fully bissecting incision was made immediately posterior to the second string. The anterior portion of the animal was placed in methacarn (6:3:1 methanol:chloroform:acetic acid) (70). Tissues in methacarn were stored at 4’C with rocking. Regular fixative changes were made when the fixative was no longer clear. Tissues were dissected and a final fixative change was performed with fresh anhydrous methacarn before incubating for one additional week, 4’C with rocking. The anhydrous methacarn was replaced with anhydrous methanol and the tissues stored at 4’C until embedding. Tissues were cleared with a decreasing methanol:xylene series then embedded in Paraplast Plus® (Millipore Sigma, St. Louis, MO U.S.A.).

### Fluorescence *In-Situ* Hybridization (FISH)

Paraffin-embedded tissues were sectioned at 6-8 *μ*m using a microtome and placed on poly-L-lysine-coated slides. Sections on slides were cleared with xylene then rehydrated with an ethanol:water series. Slides were bleached for 8-12 h using 2% hydrogen peroxide and a standard fluorescent light bulb (with concomitant cooling over an ice bath). 0.54M NaCl, 12mM Tris-Cl, 30% formamide, and 1.2% sodium doecyl sulfate with 1*μ*M Cy3 probe, 1*μ*M Cy5 probe, and 3*μ*M Eub338. Slides were observed using a Nikon A1R microscope (laser wavelengths 405nm (DAPI), 488nm (Alexa488), 558nm (Cy3), 640nm (Cy5)) and images processed using ImageJ 1.51s (71).

### Bead-beating DNA Extraction

Modified version of previous protocol published by Yu et al (72). 300 *μ*L cell lysis buffer was added to sample in bead tube containing 0.1mm and 0.5mm zirconia/silica beads (BioSpec Products, OK, U.S.A.) beads. Samples were beaten for 90 sec and briefly centrifuged before transferring supernatant to clean 1.5 mL microcentrifuge tube. An additional 150 *μ*L lysis buffer was added to the bead tube and sample was beaten again 90 sec. The supernatant previously removed was returned to bead tube and incubated for 15min at 56’C with 5 sec vortexing every 4 min. The sample was briefly centrifuged before adding 100 *μ*L 10 M ammonium acetate at 4’C and vortexed 15 sec before incubating for 10 min on ice. The sample was centrifuged for 10min at 16 000 x g. The supernatant was transferred to clean 1.5mL MCF tube and to which 350 *μ*L ethanol at 4’C was added before vortexing for 10 sec and transferring the supernatant to a QIAamp Mini spin column (Qiagen Germantown, MD U.S.A.). The sample was process as recommended in the QIAamp DNA Mini Handbook and eluted with 10 mM Tris-Cl, pH 8.5. The eluted DNA concentration was measured using Qubit™ dsDNA HS Assay Kit (Thermo Fisher Scientific Carlsbad, CA U.S.A.). Stored at −20’C.

### MasterPure DNA Extraction

MasterPure complete DNA extraction without column purification into 50*μ*L TE was performed according to manufacturer’s protocol (Epicentre Madison, WI U.S.A.). No significant difference was found between the two extraction methods (PERMANOVA: p ≥ 0.23).

### Sequencing and Initial OTU Picking

Extracted DNA samples were analyzed by amplifying the V4 hypervariable region of the 16S ribosomal RNA (rRNA) gene using primers designed in (73). PCR reactions were prepared as in (58) and sequenced using an Ilumina MiSeq (Illumina San Diego, CA U.S.A.) with custom sequencing primers added to the reagent cartridge (73) and sequenced 2 × 250bp. The sequence data was deposited in the NCBI SRA under project ID PRJNA544194.

Resulting community sequences were processed using MacQiime as outlined in (58) using a GreenGenes reference library (2013-08 release). Sequences not clustered were identified using the Ribosomal Database Project (74) to the lowest possible taxonomic level.

### Bacterial Community Analysis

Resulting sequencing data was analyzed using MacQiime and R as described below. Complete coding is available in supplemental materials and via GitHub: https://github.com/joerggraflab/McClureE_Md2019/. (32, 75)

Sample Selection: Samples with 1) fewer than 10,000 reads or 2) fewer than 3 OTUs were excluded from analysis.

Positive Controls: Two types of positive control were prepared and sequenced. 1) Amplification of a dilution series of a ZymoBIOMICS Microbial Community DNA Standard (Zymo Research, Irvine, CA U.S.A.), 2) Some samples were amplified multiple times with different PCR primers and on different runs to confirm the reproducibility of the data (data not shown). For samples amplified and sequenced multiple times, the sample with the most reads was used for analysis.

Negative Controls: Two types of negative controls were prepared and sequenced. 1) Reagent controls were prepared by performing the DNA extraction procedure using the same reagents without any sample. The resulting DNA yields for these reagent controls after extraction were always below the limit of detection for the Qubit dsDNA High-Sensitivity Assay. 2) Negative-PCR controls were prepared by performing V4-specific PCR amplification on molecular biology grade water. The resulting reactions produced no bands when analyzed with the QIAxcel DNA Fast Analysis cartridge (Qiagen Germantown, MD U.S.A.).

After sequencing, each negative control contained less than 2,000 reads. Negative controls were combined and compared to identify contaminating OTUs using maxNeg and meanNeg. maxNeg was defined as the maximum count of a single OTU found in a negative control and was determined to be 379 reads. meanNeg was defined as the mean count of all OTUs when found in negative controls and was determined to be 4 reads. OTUs were first restricted by requiring that each OTU be present in at least 1 sample with a read count >= maxNeg. This resulted in a dataset consisting of 158 OTUs.

Contaminating OTUs were further identified through the use of the decontam package (76). After removal of contaminants identified by decontam OTUs, the dataset consisted of 157 OTUs, indicating that most contaminant OTUs were found at counts <= maxNeg and so were not considered as community members.

After initial trimming, all samples contained a minimum of 10,000 reads (maximum = 375,834, minimum=10,287, median=62,015).

Effect of Extraction Method: A PERMANOVA analysis using the Bray-Curtis metric was performed through the adonis function of the vegan package (77) to determine the effect of extraction method. No significant difference was found between the two extraction methods (PERMANOVA: p ≥ 0.23).

Effect of Leech Species: A PERMANOVA analysis using the Bray-Curtis metric was performed through the adonis function of the vegan package (77) to determine the effect of leech species. This analysis was performed twice 1) with taxa split into original OTUs as determined by Qiime and 2) with taxa agglomerated at the order level (using tax_glom function of the phyloseq (75) package).

Non-metric Multidimensional Scaling (NMDS) plots were prepared using the distance, ordinate, and plot_ordination function of the phyloseq package (75). Distances were calculated with the Unifrac method (78).

Variation within *H. verbana* and *M. decora* was evaluated separately with a PERMANOVA analysis using the Bray-Curtis metric through the adonis function of the vegan package (77). Sample groups were probed for the effect of sampled organ, feeding, animal supplier/source, and shipment/collection date.

Core and Common OTUs: The average read count of a single OTU in any negative control was calculated to be 4 (meanNeg). This number was then used to calculate a conservative estimate for the minimum fraction of a sample that an OTU must compose to be considered present in that sample. It was assumed that any true OTU would contain a read count greater than or equal to meanNeg, so in the smallest sample with 10,287 total reads, an OTU must contain at least 4 reads or .03% of the total community in order to be considered present. The Core community was defined to consist of only those OTUs present in at least 90% of samples in a group. The Common community was defined to consist of only those OTUs present in at least 70% of samples in a group.

Alpha Diversity: Alpha diversity was calculated using the Shannon metric in the plot_richness function of the phyloseq package (75). Alpha diversity was calculated for ILF and intestinum samples separately.

Effect of *M. decora* Collection Month: A PERMANOVA analysis using the Bray-Curtis metric was performed through the adonis function of the vegan package (77) to determine the effect of collection month/season on *M. decora* samples. This analysis was performed twice 1) with ILF samples and 2) with intestinum samples.

NMDS plots were prepared using the distance, ordinate, and plot_ordination function of the phyloseq package (75). Distances were calculated with the Unifrac method (78).

Plotting gut community over time: For each time point after feeding, taxa were assessed for presence (≥ 0.1% of total reads) or absence. At any time point, if the taxon was determined to be present in ≤ 1 samples, it was considered absent from that time point. Ggplot2 was used to produce a stat_boxplot with whiskers at 1.5x interquartile range.

## ACKNOWLEDGEMENTS

This research was supported by NIH R01 GM095390 to J. Graf, P. Visscher, and H. Morrison, NSF 1710511 to J. Graf and V. Cooper and the University of Connecticut. We would like to thank R. Rubinstein for help with Python code used to prepare tables and the Microbial Resources and Services Facility of the University of Connecticut for sequencing the samples. J. Graf is a leech microbiology consultant for the German leech farm Biebertaler Blutegelzucht GmbH, Biebertal, Germany, and the company does not direct or approve J. Graf’s research and publications.

---

**Supplemental Table 1.** Number of samples collected of each sample type for this study. Only one sample of each sample type was collected per animal, however multiple sample types were often collected from the same animal. A total of 36 *Hirudo verbana* and 52 *Macrobdella decora* animals were used in this study.

|  | **ILF** |  | **Intestinum** |  | **Bladder** |  |
| --- | --- | --- | --- | --- | --- | --- |
| **Da1F\*\*** | ***M. decora*** | ***H. verbana*** | ***M. decora*** | ***H. verbana*** | ***M. decora*** | ***H. verbana*** |
| 0 | 17 | 10 | 15 | 2 | 8 | 2 |
| 1 | 6 | – | 6 | – | 2 | – |
| 2 | 2 | 6 | 2 | – | – | 1 |
| 4 | 6 | 4 | 9 | – | 4 | – |
| 7 | 9 | 8 | 9 | – | 3 | – |
| 30 | 7 | – | 4 | – | 2 | – |
| 90+ | 5 | 8 | – | 7 | 1 | 9 |

Supplemental Table 2. See Excel

**Supplemental Table 3.**
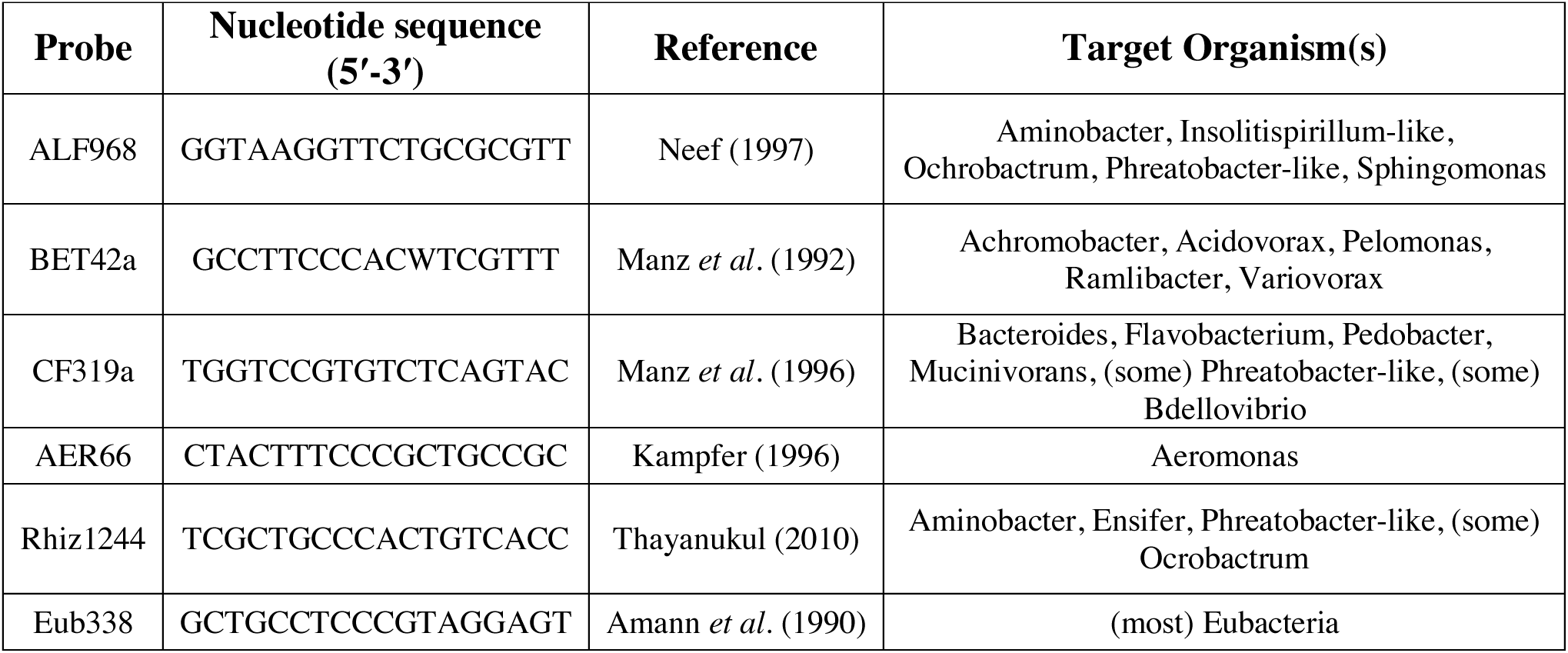
FISH probes used in this study

Sup Figure 1. Unifrac-calculated NMDS plot of wild-caught *Macrobdella decora -* associated microbiota. Animals were collected from four states of the Northeastern U.S.A.: Connecticut (orange), Massachusetts (cyan), New York (green), and Vermont (black). A) ILF samples show no significant difference between ILF of *M. decora* from CT, MA, NY, and VT (p ≥ 0.222). B) Intestinum samples from CT and MA are significantly different from each other (p = 0.006). Ellipses drawn at 95% confidence interval.

## REFERENCES

1. Sawyer RT. 1986. Leech biology and behavior. Clarendon Press, Oxford.

2. Brinkmann A, Jr. 1947. Two new antarctic leeches. Nature 160:756.

3. Whitaker IS, Rao J, Izadi D, Butler PE. 2004. Historical Article: *Hirudo medicinalis*: ancient origins of, and trends in the use of medicinal leeches throughout history. Br J Oral Maxillofac Surg 42:133–7.

4. Graf J. 2000. The symbiosis of *Aeromonas* and *Hirudo medicinalis*, the medicinal leech. ASM News 66:147–153.

5. Siddall ME, Trontelj P, Utevsky SY, Nkamany M, Macdonald KS. 2007. Diverse molecular data demonstrate that commercially available medicinal leeches are not *Hirudo medicinalis*. Proc Biol Sci 274:1481–1487.

6. Utevsky SY, Trontelj P. 2005. A new species of the medicinal leech (Oligochaeta, Hirudinida, Hirudo) from Transcaucasia and an identification key for the genus Hirudo. Parasitol Res 98:61–6.

7. Marden JN, McClure EA, Beka L, Graf J. 2016. Host Matters: Medicinal Leech Digestive-Tract Symbionts and Their Pathogenic Potential. Front Microbiol 7:1569.

8. Siddall ME, Min GS, Fontanella FM, Phillips AJ, Watson SC. 2011. Bacterial symbiont and salivary peptide evolution in the context of leech phylogeny. Parasitology 138:1815–27.

9. Rio RVM, Attardo GM, Weiss BL. 2016. Grandeur Alliances: Symbiont Metabolic Integration and Obligate Arthropod Hematophagy. Trends in Parasitology 32:739–749.

10. Min GS, Sarkar IN, Siddall ME. 2010. Salivary transcriptome of the North American medicinal leech, *Macrobdella decora*. J Parasitol 96:1211–21.

11. Whitaker IS, Oboumarzouk O, Rozen WM, Naderi N, Balasubramanian SP, Azzopardi EA, Kon M. 2012. The efficacy of medicinal leeches in plastic and reconstructive surgery: a systematic review of 277 reported clinical cases. Microsurgery 32:240–50.

12. Wenning A, Zerbst-Boroffka I, Bazin B. 1980. Water and salt excretion in the leech. J Comp Physiol B 139:97–102.

13. Graf J. 1999. Symbiosis of *Aeromonas veronii* biovar sobria and *Hirudo medicinalis*, the medicinal leech: a novel model for digestive tract associations. Infect Immun 67:1–7.

14. Worthen PL, Gode CJ, Graf J. 2006. Culture-independent characterization of the digestive-tract microbiota of the medicinal leech reveals a tripartite symbiosis. Appl Environ Microbiol 72:4775–4781.

15. Kikuchi Y, Graf J. 2007. Spatial and temporal population dynamics of a naturally occurring two-species microbial community inside the digestive tract of the medicinal leech. Appl Environ Microbiol 73:1984–1991.

16. Nelson M, Bomar L, Maltz M, Graf J. 2015. *Mucinivorans hirudinis* gen. nov., sp. nov., an anaerobic, mucin-degrading bacterium isolated from the digestive tract of the medicinal leech, *Hirudo verbana*. Int J Syst Evol Microbiol 65:990–5.

17. Neupane S, Modry D, Pafčo B, Zurek L. 2019. Bacterial Community of the Digestive Tract of the European Medicinal Leech (*Hirudo verbana*) from the Danube River. Microb Ecol doi:10.1007/s00248-019-01349-z.

18. Beka L, Fullmer MS, Colston SM, Nelson MC, Talagrand-Reboul E, Walker P, Ford B, Whitaker IS, Lamy B, Gogarten JP, Graf J. 2018. Low-Level Antimicrobials in the Medicinal Leech Select for Resistant Pathogens That Spread to Patients. mBio 9.

19. Laufer AS, Siddall ME, Graf J. 2008. Characterization of the digestive-tract microbiota of *Hirudo orientalis*, a european medicinal leech. Appl Environ Microbiol 74:6151–4.

20. Whitaker IS, Maltz M, Siddall ME, Graf J. 2014. Characterization of the digestive tract microbiota of *Hirudo orientalis* (medicinal leech) and antibiotic resistance profile. Plast Reconstr Surg 133:408e–418e.

21. Maltz MA, Bomar L, Lapierre P, Morrison HG, McClure EA, Sogin ML, Graf J. 2014. Metagenomic analysis of the medicinal leech gut microbiota. Front Microbiol 5:151.

22. Nogge G. 1981. Significance of symbionts for the maintenance of an optimal nutritional state for successful reproduction in hematophagous arthropods. Parasitology 82:101–104.

23. Manzano-Marin A, Oceguera-Figueroa A, Latorre A, Jimenez-Garcia LF, Moya A. 2015. Solving a bloody mess: B-vitamin independent metabolic convergence among gammaproteobacterial obligate endosymbionts from blood-feeding arthropods and the leech *Haementeria officinalis*. Genome Biol Evol 7:2871–84.

24. Rio RVM, Anderegg M, Graf J. 2007. Characterization of a catalase gene from *Aeromonas veronii*, the digestive-tract symbiont of the medicinal leech. Microbiol 153:1897–1906.

25. Husnik F. 2018. Host–symbiont–pathogen interactions in blood-feeding parasites: nutrition, immune cross-talk and gene exchange. Parasitology 145:1294–1303.

26. Silver AC, Graf J. 2011. Innate and procured immunity inside the digestive tract of the medicinal leech. Inv Survival Journal 8:173–178.

27. Maltz M, Graf J. 2011. The type II secretion system is essential for erythrocyte lysis and gut colonization by the leech digestive tract symbiont *Aeromonas veronii*. Appl Environ Microbiol 77:597–603.

28. Davidson SK, Powell R, James S. 2013. A global survey of the bacteria within earthworm nephridia. Mol Phylogenet Evol 67:188–200.

29. Viana F, Paz LC, Methling K, Damgaard CF, Lalk M, Schramm A, Lund MB. 2018. Distinct effects of the nephridial symbionts *Verminephrobacter* and Candidatus *Nephrothrix* on reproduction and maturation of its earthworm host *Eisenia andrei*. FEMS Microbiol Ecol 94.

30. Kikuchi Y, Bomar L, Graf J. 2009. Stratified bacterial community in the bladder of the medicinal leech, *Hirudo verbana*. Environ Microbiol 11:2758–70.

31. Siddall ME, Worthen PL, Johnson M, Graf J. 2007. Novel role for *Aeromonas jandaei* as a digestive tract symbiont of the North American medicinal leech. Appl Environ Microbiol 73:655–8.

32. Caporaso JG, Kuczynski J, Stombaugh J, Bittinger K, Bushman FD, Costello EK, Fierer N, Peña AG, Goodrich JK, Gordon JI, Huttley GA, Kelley ST, Knights D, Koenig JE, Ley RE, Lozupone CA, McDonald D, Muegge BD, Pirrung M, Reeder J, Sevinsky JR, Turnbaugh PJ, Walters WA, Widmann J, Yatsunenko T, Zaneveld J, Knight R. 2010. QIIME allows analysis of high-throughput community sequencing data. Nat Methods 7:335–336.

33. Sartor C, Bornet C, Guinard D, Fournier PE. 2013. Transmission of *Aeromonas hydrophila* by leeches. Lancet 381:1686.

34. Ge H, Jensen PD, Batstone DJ. 2011. Temperature phased anaerobic digestion increases apparent hydrolysis rate for waste activated sludge. Water Res 45:1597–1606.

35. Ott BM, Rickards A, Gehrke L, Rio RV. 2014. Characterization of shed medicinal leech mucus reveals a diverse microbiota. Front Microbiol 5:757.

36. Alexiev A, Oliverio AM, Prest TL, Korpita TM, McKenzie VJ, Song SJ, Di Fiore A, Seguin-Orlando A, Feh C, Mendelson JR, Sanders J, Knight R, Delsuc F, Amato KR, Metcalf JL, Orlando L, Kowalewski M, Avenant NL, Link A. 2017. The Effects of Captivity on the Mammalian Gut Microbiome. Integr Comp Biol 57:690–704.

37. Nelson MC, Morrison HG, Benjamino J, Grim SL, Graf J. 2014. Analysis, optimization and verification of Illumina-generated 16S rRNA gene amplicon surveys. PLoS ONE 9:e94249.

38. Pruesse E, Quast C, Knittel K, Fuchs BM, Ludwig W, Peplies J, Glöckner FO. 2007. SILVA: a comprehensive online resource for quality checked and aligned ribosomal RNA sequence data compatible with ARB. Nucleic Acids Res 35:7188–7196.

39. Kohl KD, Dearing MD. 2014. Wild-caught rodents retain a majority of their natural gut microbiota upon entrance into captivity. Environ Microbiol Reports 6:191–5.

40. Silver AC, Kikuchi Y, Fadl AA, Sha J, Chopra AK, Graf J. 2007. Interaction between innate immune cells and a bacterial type III secretion system in mutualistic and pathogenic associations. Proc Natl Acad Sci USA 104:9481–9486.

41. Clark MA, Moran NA, Baumann P, Wernegreen JJ. 2000. Cospeciation between bacterial endosymbionts (*Buchnera*) and a recent radiation of aphids (*Uroleucon*) and pitfalls of testing for phylogenetic congruence. Evolution 54:517–25.

42. Moeller AH, Li Y, Mpoudi Ngole E, Ahuka-Mundeke S, Lonsdorf EV, Pusey AE, Peeters M, Hahn BH, Ochman H. 2014. Rapid changes in the gut microbiome during human evolution. Proc Natl Acad Sci USA doi:10.1073/pnas.1419136111:201419136.

43. Chandler JA, Morgan Lang J, Bhatnagar S, Eisen JA, Kopp A. 2011. Bacterial Communities of Diverse *Drosophila* Species: Ecological Context of a Host–Microbe Model System. PLOS Genetics 7:e1002272.

44. Brune A, Dietrich C. 2015. The Gut Microbiota of Termites: Digesting the Diversity in the Light of Ecology and Evolution. Annu Rev Microbiol 69:145–166.

45. Roeselers G, Mittge EK, Stephens WZ, Parichy DM, Cavanaugh CM, Guillemin K, Rawls JF. 2011. Evidence for a core gut microbiota in the zebrafish. ISME J 5:1595–608.

46. Wang Y, Gilbreath TM, Kukutla P, Yan G, Xu J. 2011. Dynamic gut microbiome across life history of the malaria mosquito *Anopheles gambiae* in Kenya. PLoS ONE 6:e24767.

47. Aksoy S. 2000. Tsetse--A haven for microorganisms. Parasitol Today 16:114–8.

48. Strand MR. 2018. Composition and functional roles of the gut microbiota in mosquitoes. Curr Opin Insect Sci 28:59–65.

49. Zepeda Mendoza ML, Xiong Z, Escalera-Zamudio M, Runge AK, Thézé J, Streicker D, Frank HK, Loza-Rubio E, Liu S, Ryder OA, Samaniego Castruita JA, Katzourakis A, Pacheco G, Taboada B, Löber U, Pybus OG, Li Y, Rojas-Anaya E, Bohmann K, Carmona Baez A, Arias CF, Liu S, Greenwood AD, Bertelsen MF, White NE, Bunce M, Zhang G, Sicheritz-Pontén T, Gilbert MPT. 2018. Hologenomic adaptations underlying the evolution of sanguivory in the common vampire bat. Nat Ecol Evol 2:659–668.

50. Michel AJ, Ward LM, Goffredi SK, Dawson KS, Baldassarre DT, Brenner A, Gotanda KM, McCormack JE, Mullin SW, O’Neill A, Tender GS, Uy JAC, Yu K, Orphan VJ, Chaves JA. 2018. The gut of the finch: uniqueness of the gut microbiome of the Galápagos vampire finch. Microbiome 6:167–167.

51. Siddall ME, Barkdull M, Tessler M, Brugler MR, Borda E, Hekkala E. 2019. Ideating iDNA: Lessons and limitations from leeches in legacy collections. PLOS ONE 14:e0212226.

52. Colston SM, Fullmer MS, Beka L, Lamy B, Gogarten JP, Graf J. 2014. Bioinformatic genome comparisons for taxonomic and phylogenetic assignments using *Aeromonas* as a test case. mBio 5:e02136.

53. Morandi A, Zhaxybayeva O, Gogarten JP, Graf J. 2005. Evolutionary and Diagnostic Implications of Intragenomic Heterogeneity in the 16S rRNA Gene in *Aeromonas* Strains. J Bacteriol 187:6561–6564.

54. Silver AC, Rabinowitz NM, Kuffer S, Graf J. 2007. Identification of *Aeromonas veronii* genes required for colonization of the medicinal leech, *Hirudo verbana*. J Bacteriol 189:6763–72.

55. Bomar L, Graf J. 2012. Investigation into the physiologies of *Aeromonas veronii in vitro* and inside the digestive tract of the medicinal leech using RNA-seq. Biol Bull 223:155–66.

56. Maltz M, LeVarge B, Graf J. 2015. Identification of iron and heme utilization genes in *Aeromonas* and their role in the colonization of the leech digestive tract. Front Microbiol 6:763.

57. Sogin ML, Morrison HG, Huber JA, Welch DM, Huse SM, Neal PR, Arrieta JM, Herndl GJ. 2006. Microbial diversity in the deep sea and the underexplored “rare biosphere”. Proc Natl Acad Sci USA 103:12115–12120.

58. Benjamino J, Lincoln S, Srivastava R, Graf J. 2018. Low-abundant bacteria drive compositional changes in the gut microbiota after dietary alteration. Microbiome 6:86.

59. Jousset A, Bienhold C, Chatzinotas A, Gallien L, Gobet A, Kurm V, Küsel K, Rillig MC, Rivett DW, Salles JF, van der Heijden MGA, Youssef NH, Zhang X, Wei Z, Hol WHG. 2017. Where less may be more: how the rare biosphere pulls ecosystems strings. The ISME Journal 11:853–862.

60. Meyer KM, Memiaghe H, Korte L, Kenfack D, Alonso A, Bohannan BJM. 2018. Why do microbes exhibit weak biogeographic patterns? The ISME Journal 12:1404–1413.

61. Woodcock S, Curtis TP, Head IM, Lunn M, Sloan WT. 2006. Taxa-area relationships for microbes: the unsampled and the unseen. Ecol Lett 9:805–12.

62. Indergand S, Graf J. 2000. Ingested blood contributes to the specificity of the symbiosis of *Aeromonas veronii* biovar sobria and *Hirudo medicinalis*, the medicinal leech. Appl Environ Microbiol 66:4735–4741.

63. Vasai F, Brugirard Ricaud K, Bernadet MD, Cauquil L, Bouchez O, Combes S, Davail S. 2014. Overfeeding and genetics affect the composition of intestinal microbiota in *Anas platyrhynchos* (Pekin) and *Cairina moschata* (Muscovy) ducks. FEMS Microbiol Ecol 87:204–16.

64. Costello EK, Gordon JI, Secor SM, Knight R. 2010. Postprandial remodeling of the gut microbiota in Burmese pythons. The ISME journal 4:1375–1385.

65. Tasiemski A, Massol F, Cuvillier-Hot V, Boidin-Wichlacz C, Roger E, Rodet F, Fournier I, Thomas F, Salzet M. 2015. Reciprocal immune benefit based on complementary production of antibiotics by the leech *Hirudo verbana* and its gut symbiont *Aeromonas veronii*. Sci Rep 5:17498.

66. Carey HV, Walters WA, Knight R. 2013. Seasonal restructuring of the ground squirrel gut microbiota over the annual hibernation cycle. Am J Physiol Regul Integr Comp Physiol 304:R33–R42.

67. Ott BM, Beka L, Graf J, Rio RV. 2015. Draft Genome Sequence of *Pedobacter* sp. Strain Hv1, an Isolate from Medicinal Leech Mucosal Castings. Genome Announc 3.

68. Silver AC, Williams D, Faucher J, Horneman AJ, Gogarten JP, Graf J. 2011. Complex evolutionary history of the *Aeromonas veronii* group revealed by host interaction and DNA sequence data. PLoS One 6:e16751.

69. Mumcuoglu KY, Huberman L, Cohen R, Temper V, Adler A, Galun R, Block C. 2010. Elimination of symbiotic *Aeromonas* spp. from the intestinal tract of the medicinal leech, *Hirudo medicinalis*, using ciprofloxacin feeding. Clin Microbiol Infect 16:563–7.

70. Puchtler H, Waldrop FS, Meloan SN, Terry MS, Conner HM. 1970. Methacarn (methanol-Carnoy) fixation. Practical and theoretical considerations. Histochemie 21:97–116.

71. Schneider CA, Rasband WS, Eliceiri KW. 2012. NIH Image to ImageJ: 25 years of image analysis. Nat Methods 9:671–675.

72. Yu Z, Morrison M. 2004. Improved extraction of PCR-quality community DNA from digesta and fecal samples. Biotechniques 36:808–12.

73. Kozich JJ, Westcott SL, Baxter NT, Highlander SK, Schloss PD. 2013. Development of a Dual-Index Sequencing Strategy and Curation Pipeline for Analyzing Amplicon Sequence Data on the MiSeq Illumina Sequencing Platform. Appl Environ Microbiol 79:5112–5120.

74. Larsbrink J, Rogers TE, Hemsworth GR, McKee LS, Tauzin AS, Spadiut O, Klinter S, Pudlo NA, Urs K, Koropatkin NM, Creagh AL, Haynes CA, Kelly AG, Cederholm SN, Davies GJ, Martens EC, Brumer H. 2014. A discrete genetic locus confers xyloglucan metabolism in select human gut Bacteroidetes. Nature doi:10.1038/nature12907.

75. McMurdie PJ, Holmes S. 2013. phyloseq: An R Package for Reproducible Interactive Analysis and Graphics of Microbiome Census Data. PLOS ONE 8:e61217.

76. Davis NM, Proctor DM, Holmes SP, Relman DA, Callahan BJ. 2018. Simple statistical identification and removal of contaminant sequences in marker-gene and metagenomics data. Microbiome 6:226.

77. Oksanen J, Blanchet FG, Friendly M, Kindt R, Legendre P, McGlinn D, Minchin PR, O’Hara RB, Simpson GL, Solymos P, Stevens MHH, Szoecs E, Wagner H. 2019. vegan: Community Ecology Package. R package version 2.5–5, https://CRAN.R-project.org/package=vegan.

78. Lozupone C, Knight R. 2005. UniFrac: a New Phylogenetic Method for Comparing Microbial Communities. Appl Environ Microbiol 71:8228–8235.

